# Basolateral amygdala dopamine transmits emotional salience

**DOI:** 10.1101/2025.05.15.654323

**Authors:** Megan A. Brickner, William E. Szot, Amy R. Wolff, Mark J. Thomas, Benjamin T. Saunders

**Affiliations:** Department of Neuroscience, University of Minnesota; Medical Discovery Team on Addiction, University of Minnesota; Graduate Program in Neuroscience, University of Minnesota

## Abstract

Adaptive decision making relies on proper disambiguation of upcoming positive and negative events. The basolateral amygdala (BLA) is central to this, assigning emotional value to stimuli to facilitate appropriate behavior. The ventral tegmental area (VTA), which is classically known to regulate associative learning and incentive motivation via dopamine projections to the striatum, also contains strong dopamine projections to the BLA, but this system has received much less attention. Here, we investigated how in vivo BLA dopamine signaling is engaged during valenced experiences in associative learning. First, we show that BLA dopamine signals scale with intensity of aversive stimuli, but not reward value, and reward-predictive cues evoke BLA dopamine signals that diminish, rather than grow, with training. As the complexity of the learning context was increased, where rats actively differentiated between various cue types signaling threat, reward, safety, and neutral associations, the magnitude of cue-evoked BLA dopamine re-scaled to report the level of relative emotional saliency. Fear and safety cues prompted larger, more sustained dopamine signals compared to reward and neutral cues. Despite these distinctions, cue-evoked dopamine signals did not reliably correlate with behavioral expression, and optogenetically enhancing BLA dopamine did not alter freezing behavior. Together, these results indicate that dopamine signaling in the BLA encodes the emotional weight of sensory transitions, but not the associative strength or value of conditioned cues. Our findings broaden the theoretical landscape of dopamine heterogeneity, showing that BLA dopamine contributes to tagging affective states, in support of dynamic discernment of relative stimulus importance during learning.

## INTRODUCTION

A foundation of behavior is the associative learning that cements relationships between environmental cues and biologically relevant outcomes. Disambiguation of positive and negative valenced sensory inputs is then imperative for directing appropriate decision making. The basolateral amygdala (BLA) is a key brain structure in this process, integrating sensory information from multiple brain regions and assigning emotional value to environmental cues and contexts (Paton et al., 2006; O’Neill et al., 2018). The BLA is central to fear (Maren, 2005; Duvarci and Pare, 2014) as well as reward learning (Cador et al., 1989; Everitt et al., 1991; Whitelaw et al., 1996; Baxter and Murray, 2002; Ambroggi et al., 2008; Wassum and Izquierdo, 2015; Costa et al., 2016; Sias et al., 2021; Ottenheimer et al., 2024), and is involved in discrimination of fear and reward (Sangha et al., 2013, 2020; Duvarci and Pare, 2014; Janak and Tye, 2015; Sangha, 2015; Ng et al., 2018; Sias et al., 2024) via dynamic valence encoding and memory representation (Rogan et al., 2005; Tye et al., 2008; Sangha et al., 2013; Josselyn et al., 2015; Sengupta et al., 2018).

The BLA’s role in emotional learning is partly driven by excitatory inputs from cortex, thalamus and other regions, but the contribution of neuromodulatory circuit inputs is less clear (Saddoris et al., 2005; Paton et al., 2006; Namburi et al., 2015; McGarry and Carter, 2017; Beyeler et al., 2018; O’Neill et al., 2018; Likhtik and Johansen, 2019; Khalil et al., 2023; Leppla et al., 2023; Li et al., 2025). Notably, the ventral tegmental area (VTA), where dopamine neurons are classically known to regulate reward learning and incentive motivation via projections to the striatum (Fields et al., 2007; Yang et al., 2018) also contains dense dopamine projections to the BLA (Haber and Fudge, 1997; Brinley-Reed and McDonald, 1999; Takahashi et al., 2010; Mingote et al., 2015; Lutas et al., 2019), and this system has received much less attention. On slow timescales, amygdala dopamine levels track discrimination learning (Hori et al., 1993). Recent studies offer more insights, showing that the dopaminergic VTA-BLA system is rapidly activated during rewards and punishments (Correia and Goosens, 2016; Lutas et al., 2019, 2022; Tang et al., 2020; Sias et al., 2024) and responds to predictive cues (Correia and Goosens, 2016; Lutas et al., 2019; Tang et al., 2020;

Grove et al., 2022; Morel et al., 2022; Sias et al., 2024). Further, BLA dopamine signaling encodes information relating to reward-identity memories (Correia and Goosens, 2016; Lutas et al., 2019; Tang et al., 2020; Sias et al., 2024). Despite this growing focus, we still know very little about how the BLA dopamine system is engaged during learning. Delineating this aspect of heterogeneity is critical for understanding evolving complexities of the role of dopamine in valuation and salience encoding (McCutcheon et al., 2012; Lammel et al., 2014; Kutlu et al., 2021; van Elzelingen et al., 2022a, 2022b; Bornhoft et al., 2025).

To investigate this, we used fiber photometry and optogenetics to characterize in vivo BLA dopamine signals as rats engaged in various learning tasks. We first show that appetitive and aversive stimuli evoke strong positive dopamine responses in the BLA, and reward-cue evoked responses diminish, rather than grow, with training. We then tracked the profile of BLA dopamine in response to four differently valenced cue types during a Pavlovian cue discrimination task. In this more complex learning context, where cues signaling threat, reward, safety, and neutral associations were intermingled within the same session, we found that the magnitude of cue-evoked BLA dopamine responses scaled with relative emotional saliency: fear and safety cues evoked strong signals early in conditioning that were larger than for reward and neutral cues, and these signals were resistant to extinction. In additional studies, we demonstrate optogenetic stimulation of dopamine release in the BLA does not directly drive learning via reinforcement, and BLA dopamine signals do not track the value of preferred rewards, but instead scale with stimulus uncertainty and threat intensity. Together, our data reveal a new pattern of dopamine signaling with a unique contribution to learning compared to classic striatal systems, through encoding the emotional weight of sensory state transitions, independent of associative strength or value. This state-guided affective signal is dynamically engaged when relative stimulus importance must be disambiguated.

## RESULTS

To record amygdala dopamine signaling in freely behaving rats, we used fiber photometry with the dLight1.3b sensor (**Fig 1A**). First, to validate this approach and determine how VTA TH-neuron stimulation influences dopamine dynamics in the BLA, a cre-driven Chrimson opsin was expressed into the VTA of TH-cre rats to specifically target the dopamine neurons (**Fig 1B,D**), while dLight1.3b was expressed in the ipsilateral BLA (**Fig 1B,C; Fig S1**). Optogenetic activation of VTA dopamine neurons via 590-nm laser pulses evoked robust dopa-mine signals in the BLA, which positively scaled with stimulation frequency (**Fig 1E,F**; t(7)=5.031, p=0.0015). We next targeted optogenetic activation to VTA dopamine terminals in the BLA to confirm locally evoked dopamine release. Terminal stimulation also resulted in robust dopamine signaling (**Fig 1G**) that was comparable to that evoked by brief dopamine cell body stimulation (**Fig 1H**; t(7)=1.858, p=0.1055).

**Fig 1.**
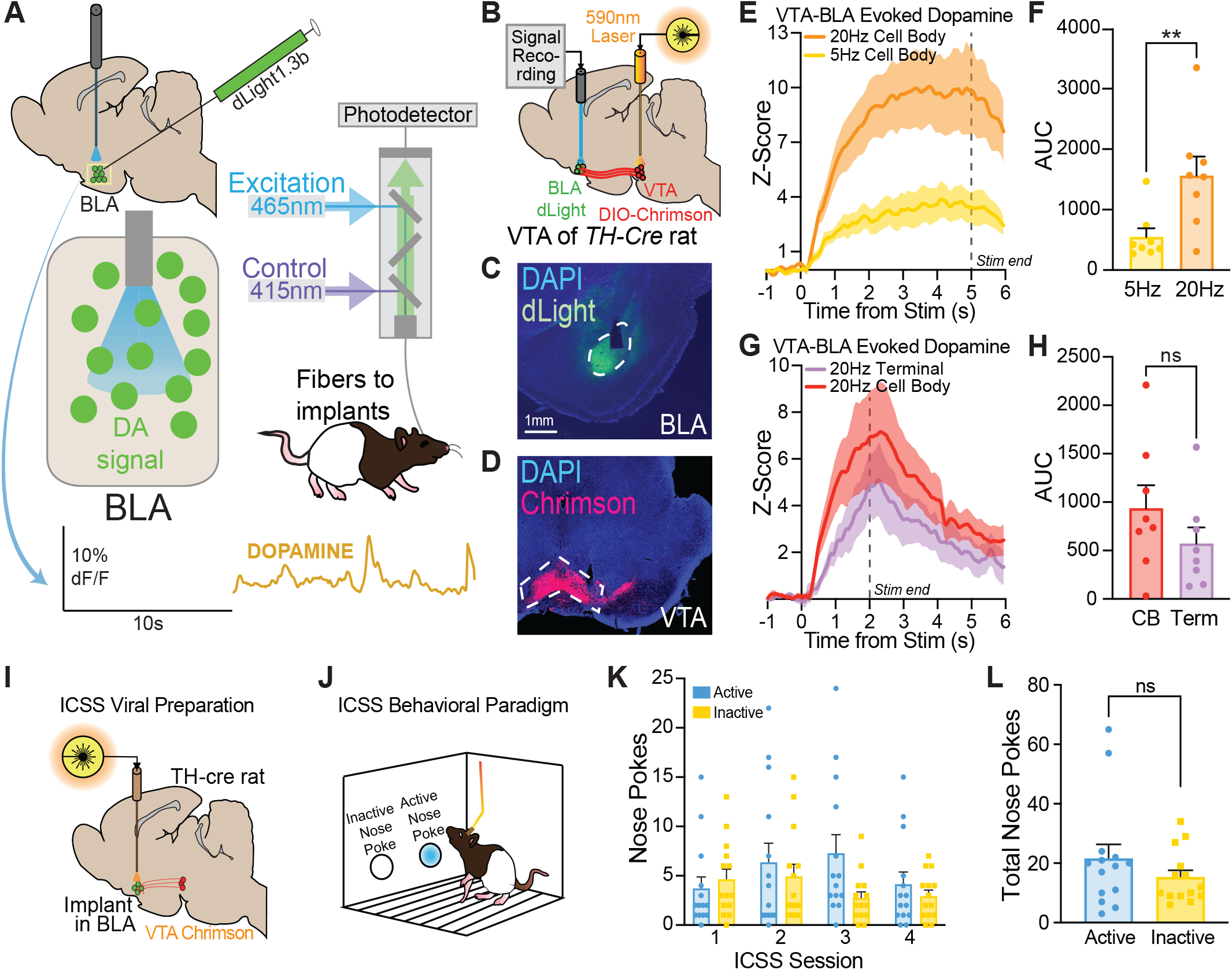
Optogenetic activation of VTA dopamine neurons evokes robust BLA dopamine signaling that does not support reinforcement. A) Schematic of the photometry system to record BLA dopamine dynamics. B) Viral approach for simultaneous in vivo optogenetic activation VTA dopamine neuron cell bodies or terminals in the BLA paired with fiber photometry recording of dopamine transmission in the BLA. Dopamine neurons were targeted in TH-cre transgenic rats with DIO-Chrimson opsin (n=8; 4F, 4M). C) Example of dLight1.3b viral expression in the BLA, with tissue counterstained for visualization of overlapping tyrosine hydroxylase (TH). D) Example of DIO-Chrimson expression in VTA dopamine neurons. E, F) Optogenetic excitation of VTA dopamine cell bodies resulted in frequency-dependent dopamine release in the BLA, where 5 s stimulation at 20Hz prompted a larger response compared to 5 s stimulation at 5Hz. G, H) Optogenetic excitation of VTA dopamine neuron terminals in the BLA also resulted in robust BLA dopamine release, which was comparable to the signal evoked by cell body stimulation for 2 s at 20Hz. I) Viral approach for intracranial self stimulation (ICSS) of VTA-BLA dopamine terminals using DIO-Chrimson or DIO-ChR infused in the VTA of TH-cre rats and optic fibers implanted over the BLA. J) Schematic of the ICSS behavioral paradigm, where rats could nose poke to activate VTA-BLA dopamine terminals to prompt dopamine transmission into the BLA. K) Rats (n=14; 6F, 8M) did not readily poke for BLA dopamine terminal activation during ICSS, making the same number of active nose pokes (ANPs) and inactive nose pokes (INPs). L) Total number of ANPs and INPs during ICSS collapsed across sessions. AUC = Area under curve. Data represent mean ± SEM. *p<0.05, **p<0.01.

VTA dopamine projections to the striatum are known to support behavioral reinforcement (Witten et al., 2011; Kim et al., 2012; Steinberg et al., 2014; Wolff and Saunders, 2024). Given the strong evoked dopamine signal we observed in the BLA, we next tested whether VTA-BLA dopamine transmission also promotes reinforcement (**Fig 1I**). Rats were given the opportunity to nose poke for optogenetic activation of VTA dopamine terminals in the BLA, in an Intracranial Self Stimulation (ICSS) procedure (**Fig 1J**). Overall, rats did not readily nose poke for stimulation (**Fig 1K**; F(1,13)=3.059, p=0.1039), and active nose pokes were not significantly different from inactive nose pokes (**Fig 1L**; t(13)=1.749, p=0.1039).

### In vivo BLA dopamine signals track stimulus intensity but not value

We next recorded BLA dopamine dynamics in response to differently valenced unconditioned stimuli - food reward or footshock. Rats were first given the opportunity to drink sucrose, delivered into a magazine port, which prompted a moderate increase in dopamine (**Fig 2A**). Delivery of moderate (0.4mA, 0.5s) footshocks elicited a strong phasic increase in dopamine (**Fig 2B**). Comparing the signals directly, footshock-evoked signals were significantly larger than sucrose evoked signals (**Fig 2C**; t(18)=4.191, p=0.0005). Broadly, these results show that both positive and negative events prompt release of BLA dopamine, with a bias of stronger responses for aversive stimuli.

**Fig 2.**
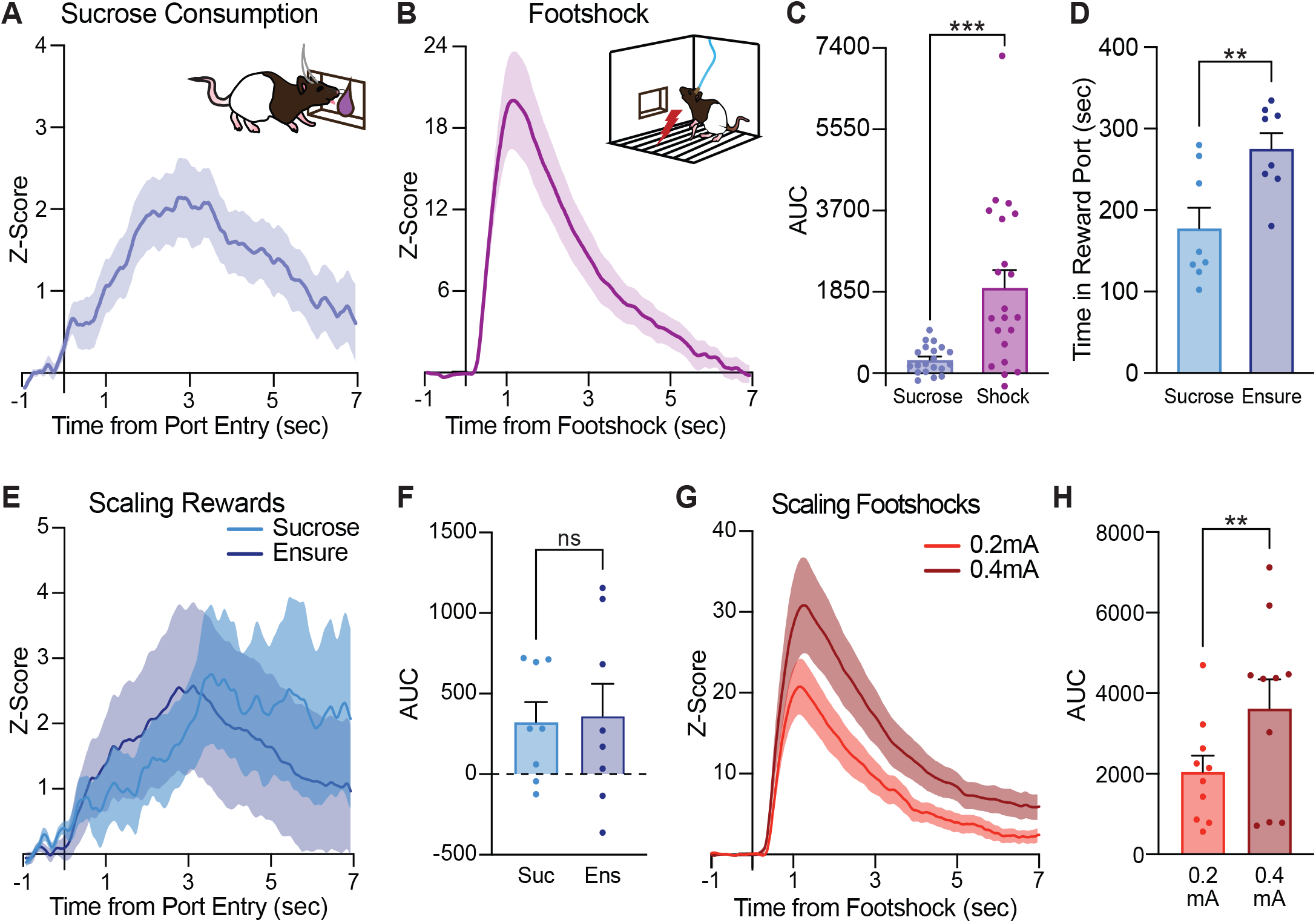
In vivo BLA dopamine signals track stimulus emotional intensity but not value. A) Rats (n=21) were given unsignalled deliveries of a sucrose reward. Sucrose consumption prompted modest increases of in vivo BLA dopamine. Photometry traces were time locked to the first reward port entry after sucrose pump onset. B) Delivery of moderate 0.4mA footshocks evoked large BLA dopamine responses in the same rats. C) The footshock-evoked signal was larger than the BLA dopamine response to sucrose consumption when comparing AUC values. D) Rats (n=8) spend significantly more time in the ensure reward port compared to the sucrose reward port. E) Sucrose consumption and ensure consumption elicit moderate increases of BLA dopamine, F) that are equal in AUC magnitude of the first 5 s after rewarded port entry onset. G) Rats (n=10) were exposed to uncued footshocks. Footshocks of both mild (0.2mA) and moderate (0.4mA) currents prompt robust BLA dopamine transients. H) BLA dopamine responses were significantly larger to the higher magnitude shocks. Data represent mean ± SEM. *p<0.05, **p<0.01, ***p<0.001.

To dissociate how BLA dopamine signals are engaged by stimuli of different values versus different intensities, we compared responses to sucrose versus a highly palatable Ensure solution. In consumption tests, rats preferred the Ensure reward, spending more time in the Ensure magazine compared to the sucrose magazine (**Fig 2D**; t(7)=3.551, p=0.0093). However, both reward types prompted modest increases in BLA dopamine (**Fig 2E**) that were equal in magnitude (**Fig 2F**; t(7)=0.2346, p=0.8212). We next tracked BLA dopamine responses to random, brief footshocks of either 0.2mA or 0.4mA (**Fig 2F**). Both footshock intensities evoked large dopamine transients, but the stronger footshock level evoked greater dopamine release in the BLA (**Fig 2F,G**; t(9)=3.345, p=0.0086). Overall, these patterns suggest that BLA dopamine release scales within relative stimulus intensity (**Fig 2F,G**), but does not scale with relative stimulus value (**Fig 2D,E**).

### Reward learning promotes diminishing cue-evoked BLA dopamine signals across training

To explore how BLA dopamine dynamics track associative learning and valence discrimination, rats were prepared for BLA dopamine photometry and then trained on a multi-phase Pavlovian cue discrimination paradigm adapted from previous studies (Sangha et al., 2013) (**Fig 3A**). Rats first underwent reward-only conditioning (RC), followed by a compound reward and fear conditioning phase (CC). This was then followed by the full cue discrimination task (CD), where rats had to distinguish between four different, interleaved Pavlovian cue associations - reward, fear, safety, and neutral. Finally, reward and fear cues underwent extinction (**Fig 3A**). Throughout this, we measured two main conditioned responses: sucrose port entries and freezing, which were largely similar across males and females (**Fig S2**). Further, in our assessment of the photometry data, we did not see any consistent sex-related biases in BLA dopamine signals nor qualitatively different patterns of activity (**Fig S3**), in various task phases. These results are interesting in light of previous data showing a wide range of sex differences in behavioral threat and safety responses (Gruene et al., 2015; Orsini et al., 2016; Yokota et al., 2017; Chowdhury et al., 2019; Greiner et al., 2019), suggesting complex and distinct fear response strategies between female and male rodents. While we found no obvious connection between cue-evoked dopamine and behavior in our studies, our data do not rule out possible sex-biased dopaminergic mechanisms of behavioral vigilance and decision making within the amygdala (Wheeler et al., 2024).

**Fig 3.**
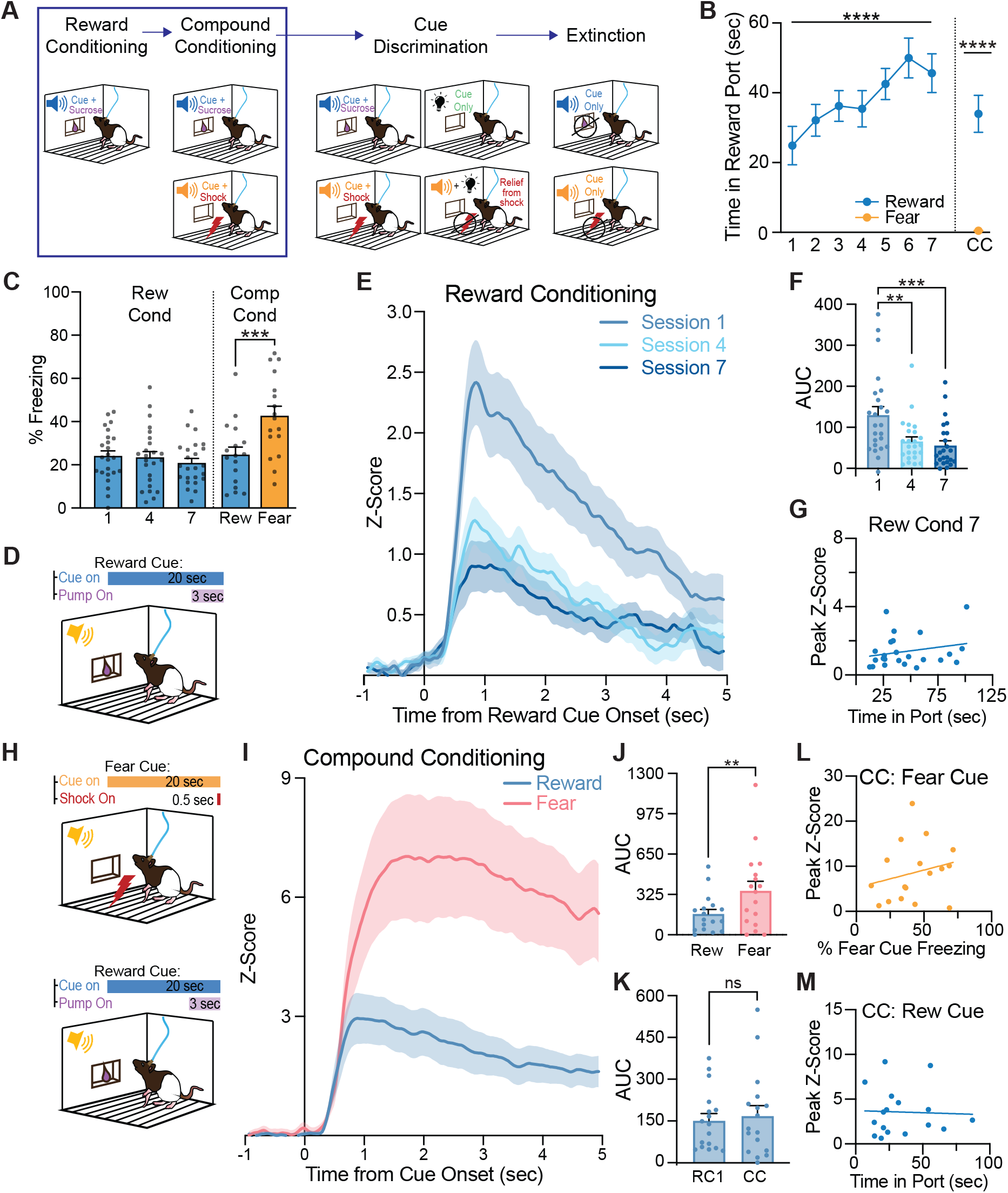
BLA dopamine signals preferentially represent conditioned threat over reward expectation. A) Schematic of the multi-phase Pavlovian cue discrimination paradigm (n=24; 11F, 13M). Behavior and photometry data for the first two phases: reward conditioning and compound (reward and fear) conditioning are shown. B) Throughout training, time spent in the reward port increased during reward cue presentations, while fear cue presentations did not prompt port entry behaviors. C) Cue-induced freezing was low for the reward cue during both reward and compound conditioning, but was higher for the fear cue in compound conditioning. D) During reward conditioning, a 20-s auditory cue was paired with a 3-s delivery of sucrose into the reward port. E) Reward cue evoked BLA dopamine responses were largest initially and decreased with training, F) as shown by AUC comparisons of the first 2 s after reward cue onset. G) There was no correlation between port entry behavior and reward cue peak dopamine signals on the final day of reward conditioning. H) During compound conditioning (n=17; 7F, 10M), trials containing a new 20 s auditory cue that predicted a footshock (0.5-s 0.5-mA) were intermingled with reward cue trials. I) Rats exhibited large dopamine dynamics to the fear cue and moderate dopamine dynamics to the reward cue. J) The fear cue-evoked signal was greater than the reward cue when comparing AUC values. K) The reward cue during both reward conditioning day 1 and compound conditioning produced similarly sized dopamine events that did not show different AUC comparisons. L) There is no correlation between conditioned freezing behaviors and the magnitude of the fear cue signal, M) or between conditioned port entry behaviors and the magnitude of the reward cue signal. Data represent mean ± SEM. *p<0.05, **p<0.01, ****p<0.0001. CC = compound conditioning. RC = reward conditioning.

In the simple reward conditioning phase, an auditory cue predicted sucrose reward delivery (**Fig 3D**). Rats learned this association quickly and increased their sucrose port entry behaviors across training (**Fig 3B**; F(6,135)=5.187, p<0.0001), while conditioned freezing behaviors were minimal (**Fig 3C**; F(2,69)=0.4699, p=0.6171). Females and males responded with similar levels of locomotion and port entry behavior during reward conditioning. While there were no significant differences in freezing during reward cue presentations for females and males, the trajectory of response patterns was different. Freezing behaviors decreased for the females, but stayed slightly elevated for the males (**Fig S2**; session x sex interaction, F(2,44)=4.566, p=0.0158). Reward cue presentations evoked dopamine increases on all days of training (**Fig 3E**). Notably, cue-evoked do-pamine was the greatest on the first day of conditioning and decreased significantly as reward conditioning progressed (**Fig 3F**; F(2,46)=10.73, p=0.0002; post hoc comparisons for session 4 vs 1, p=0.0017; session 7 vs 1, p=0.0003). There was no relationship between cue-evoked dopamine signals and the intensity of port entry behavior (**Fig 3G**; Pearson correlation, p= 0.2702).

### BLA dopamine signals preferentially represent conditioned threats over reward expectation

Following simple reward conditioning, a cue predicting footshock was added, interleaved with reward cue trials in a compound conditioning session (**Fig 3H**). On this session, rats maintained their sucrose port entry behaviors during the reward cue and spent significantly more time in the sucrose port during the reward cue compared to the fear cue (**Fig 3B**; t(16)=6.383, p<0.0001). Freezing behavior rapidly emerged to the fear cue but not the reward cue (**Fig 3C**; t(16)=4.455, p=0.0004), and reward cue freezing did not differ from simple reward conditioning (**Fig 3C**; t(16)=1.394, p=0.1824). This indicates rats were able to behaviorally discriminate the cues during compound conditioning, with similar levels of conditioned port entry behaviors and cued locomotion between females and males (**Fig S2**).

We examined BLA dopamine signals on this session, when both reward and fear learning were engaged. The fear cue elicited a significantly larger and more sustained dopamine peak than the reward cue signal (**Fig 3I,J**; t(16)=3.465, p=0.0032). Notably, on this compound conditioning session, the reward cue evoked dopamine signal rebounded relative to the final day of simple reward conditioning (**Fig 3E,I**), returning to the same magnitude it showed during the first day of reward condi-tioning (**Fig 3K**; t(16)=0.8966, p=0.3832). Despite these large signal differences for fear versus reward cues, there were no significant correlations between fear or reward cue-evoked dopamine and freezing (**Fig 3L**; Pearson correlation, p= 0.3881) or port entry behavior (**Fig 3M**; Pearson correlation, p= 0.8830).

### Rats discriminate fear, safety, and reward predictive cues

Previous in vivo single unit recordings have demonstrated that a population of BLA neurons become responsive during cue discrimination training, including to a safety cue that designates relief from shock (Sangha et al., 2013). We hypothesized that BLA dopamine transmission may be an important contributor to the disambiguation of differently valenced, salient events. Following fear plus reward compound conditioning rats were trained to distinguish between 4 different cues predicting various outcomes presented pseudorandomly during each cue discrimination behavioral session: the previously experienced (1) reward cue (paired with sucrose) and (2) fear cue (paired with a footshock), as well as a (3) neutral cue with no paired outcome and a (4) compound cue (fear+neutral cue played concurrently) to signal a relief or “safety” from shock (**Fig 4A,B**).

**Fig 4.**
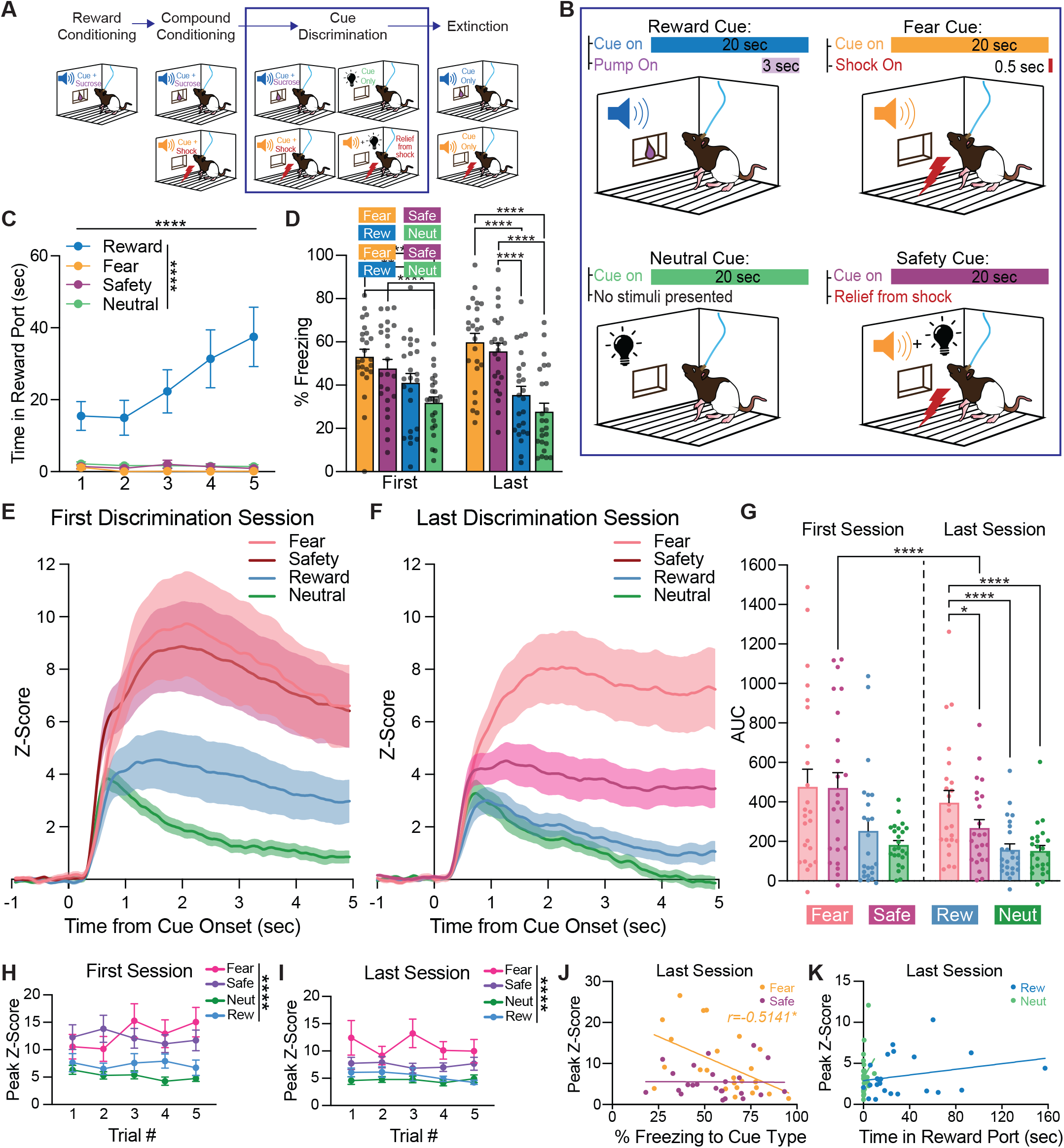
Cue-evoked BLA dopamine dynamics scale with relative emotional saliency during discrimination learning. A) Schematic of the multi-phase Pavlovian cue discrimination paradigm the rats were trained on (n= 24; 11F, 13M). Behavior and photometry data from the valenced cue discrimination phase are shown. B) During the cue discrimination task, four cues were presented randomly throughout the two hour session. The previously learned reward and fear cues were intermingled with a neutral cue (20-s visual cue with no paired behavioral outcome) and a safety cue (20 s auditory fear and visual neutral cues presented together to signify relief from shock). C) Rats spent the highest amount of time in the reward port during the reward cue, and spent almost no time in the reward port during fear, safety, or neutral cue presentations. This discrimination grew across training days as rats increased their time in port during reward cue presentations. D) Cue-induced freezing remained high for the fear and safety cue throughout discrimination training, and was significantly higher than the reward and neutral cue freezing levels by the final discrimination session. E) All four cue types elicited a positive peak on the first discrimination session, with the fear and safety cues prompting similar trace dynamics and the reward and neutral cues prompting similar trace dynamics. F) By the last discrimination session, the fear cue produced the largest dopamine trace, in terms of peak and duration, while the safety, reward, and neutral cues produced modest and transient increases in BLA dopamine. The safety cue evoked much less dopamine compared to early training. G) AUC values of the BLA dopamine signals comparing the first and last discrimination session. H) Throughout the first discrimination session, the peak values of the various dopamine traces varied based on cue type, such that by the final block of cue presentations, both the fear and safety cues elicited signals with higher peak values compared to the reward and neutral cues. I) By the last fear discrimination session, the fear cues maintained their robust and sustained BLA dopamine responses and were generally larger than the safety, reward, and neutral cue presentations. J) There was no correlation between freezing behavior and the safety cue evoked dopamine peaks, while the fear cue dopamine signal negatively correlated with fear cue-induced freezing behavior by the end of discrimination training. K) The dopamine signal for the reward and neutral cues were not correlated with port entry behaviors on the last discrimination session. Data represent mean ± SEM. *p<0.05, **p<0.01, ***p<0.001, ****p<0.0001.

During this experimental phase, rats spent more time in the sucrose port during reward cue presentation periods than any other cue period (**Fig 4C**; F(3,69)=18.24, p<0.0001). They quickly discriminated cue types and this discrimination grew across sessions (**Fig 4C**; session x cue type interaction F(12,258)=7.410, p<0.0001). Additionally, cue-induced freezing was modulated by cue type (**Fig 4D**; F(3,69)=35.46, p<0.0001), with a graded response in freezing across fear, safety, reward, and neutral cues. Further, fear cue discrimination grew across sessions (**Fig 4D**; session x cue type interaction F(3,69)=4.694, p=0.0048). Compared to the reward and neutral cues, the fear cue elicited more freezing behaviors in both the first (p=0.0025; p<0.0001) and last training session (p<0.0001; p<0.0001). The safety cue also evoked significantly more freezing than the reward cue (p<0.0001) by the last cue discrimination day and the neutral cue presentations on both the first and last cue discrimination days (p<0.0001; p<0.0001). There were no significant changes to reward or neutral cue freezing across training, which remained low. When comparing males and females, there were no significant differences in port entry behaviors to any of the cue types. There were also no sex differences in cued freezing behaviors during the cue discrimination phase. However, females exhibited significantly more locomotion than males during the neutral cue presentations (**Fig S2**).

### BLA dopamine responds preferentially to fear and safety cues throughout discrimination learning

Prominent increases in BLA dopamine were seen in response to all cues during the first (**Fig 4E**) and last (**Fig 4F**) discrimination training sessions, with different cues prompting different dopamine levels as the sessions progressed (**Fig 4G**, AUC session x cue interaction, F(3,69)=2.746, p=0.0495). The fear and safety cues evoked larger responses than the reward and neutral cues (**Fig 4E-G**), but there were no differences between the AUC values for the fear and safety signals (p=0.9993) nor the reward and neutral signals (p=0.3787) in the first discrimination session. Early in discrimination training, the dopamine responses to the fear and safety cues were characterized by large peaks followed by a sustained elevation, while the reward and neutral cues prompted smaller peaks that more readily returned to baseline (**Fig 4E; Fig S4**). In contrast, by the last discrimination session (**Fig 4F; Fig S4**), the reward and neutral cues maintained a similar evoked peak but returned to baseline more quickly. Additionally, within the last discrimination session, the fear cue elicited a similar high peak and sustained signal, but the safety cue signal diminished relative to session 1 (**Fig 4G**; AUC: p<0.0001), reaching an intermediate level interposed between the fear cue trace and the reward and neutral cue traces (**Fig 4F**). The reward cue response also decreased relative to session 1 (**Fig 4G**; AUC p=0.0325), mirroring the pattern seen during reward conditioning training (**Fig 3F**). On the last discrimination day, the fear cue presentations elicited larger and more sustained dopamine dynamics than the safety cue (p=0.0231), reward cue (p<0.0001), and neutral cue (p<0.0001; **Fig 4G**).

Throughout the first cue discrimination session, cued dopamine levels were rapidly updated as rats began to discriminate the different cue types (**Fig 4H**; trial x cue type interaction F(12,270)=1.876, p=0.0373), as demonstrated during the final block of cue presentations, where both the fear and safety cues elicited signals with higher peak values compared to reward (fear: p=0.0001; safety: p=0.0467) and neutral cues (fear: p<0.0001; safety: p=0.0018). Fear cues also continued to elicit strong and sustained BLA dopamine responses, which were generally greater than those evoked by the safety, reward, or neutral cues (**Fig 4I**; F(3,69)=15.18, p<0.0001). During the final cue discrimination session, there was no relationship between safety cue-evoked dopamine and freezing behavior (**Fig 4J**; Pearson correlation, p= 0.9615). However, after discrimination learning, but a mild negative relationship between the peak dopamine to the fear cue and conditioned freezing behavior (**Fig 4J**; Pearson correlation, p= 0.0102). There were no relationships found between evoked dopamine and conditioned port entry behaviors for the reward cue (Pearson correlation, p= 0.2234) or the neutral cue (Pearson correlation, p= 0.2918) on the last cue discrimination session (**Fig 4K**).

Overall, in various conditioning contexts, we saw positive-going in vivo BLA dopamine signals during rewarding and aversive events and their predictive cues (**Fig 2A,B,D,G; Fig 3E,H; Fig 4E,F**). Cue-evoked signals had clear qualitative distinction in the representation of different types of cues by the end of discrimination training (**Fig 4E,F**). While there was a general bias toward the more fearful or ambiguous cues, dopamine responses were larger and more preferentially biased toward threat-related cues, in particular early in discrimination learning.

### Reward cue-evoked dopamine signals dynamically rescale with changing emotional uncertainty

Presentations of a reward cue predicting a sucrose reward delivery (**Fig 5A**) reliably elicited positive BLA dopamine dynamics during the multi-phase Pavlovian cue discrimination paradigm (**Fig 3E,I; Fig 4E,F**). When comparing the conditioned behaviors during the first and last reward conditioning sessions, compound conditioning, and the first and last cue discrimination sessions, we saw that the time spent in the sucrose port varied by phase (**Fig 5B**; F(4,83)=8.149, p<0.0001). Specifically, reward seeking behaviors increased across reward conditioning (day 1 vs. day 7: p=0.0025) and cue discrimination (first vs. last day: p=0.0128), but the time spent in the reward port during reward cue presentations was blunted on the first cue discrimination session compared to the final reward conditioning session (p<0.0001). Likewise, the reward cue evoked dopamine signals across the multi-phase paradigm also varied by phase (**Fig 5C**). In general, the cued dopamine responses were characterized by a quick rise to a peak of differing magnitudes followed by varying levels of maintained activity. The dopamine signals were biased in magnitude according to phase for AUC (**Fig 5D**; F(4,85)=9.091, p<0.0001) and peak value measures (**Fig 5E**; F(4,85)=9.455, p<0.0001),

**Fig 5.**
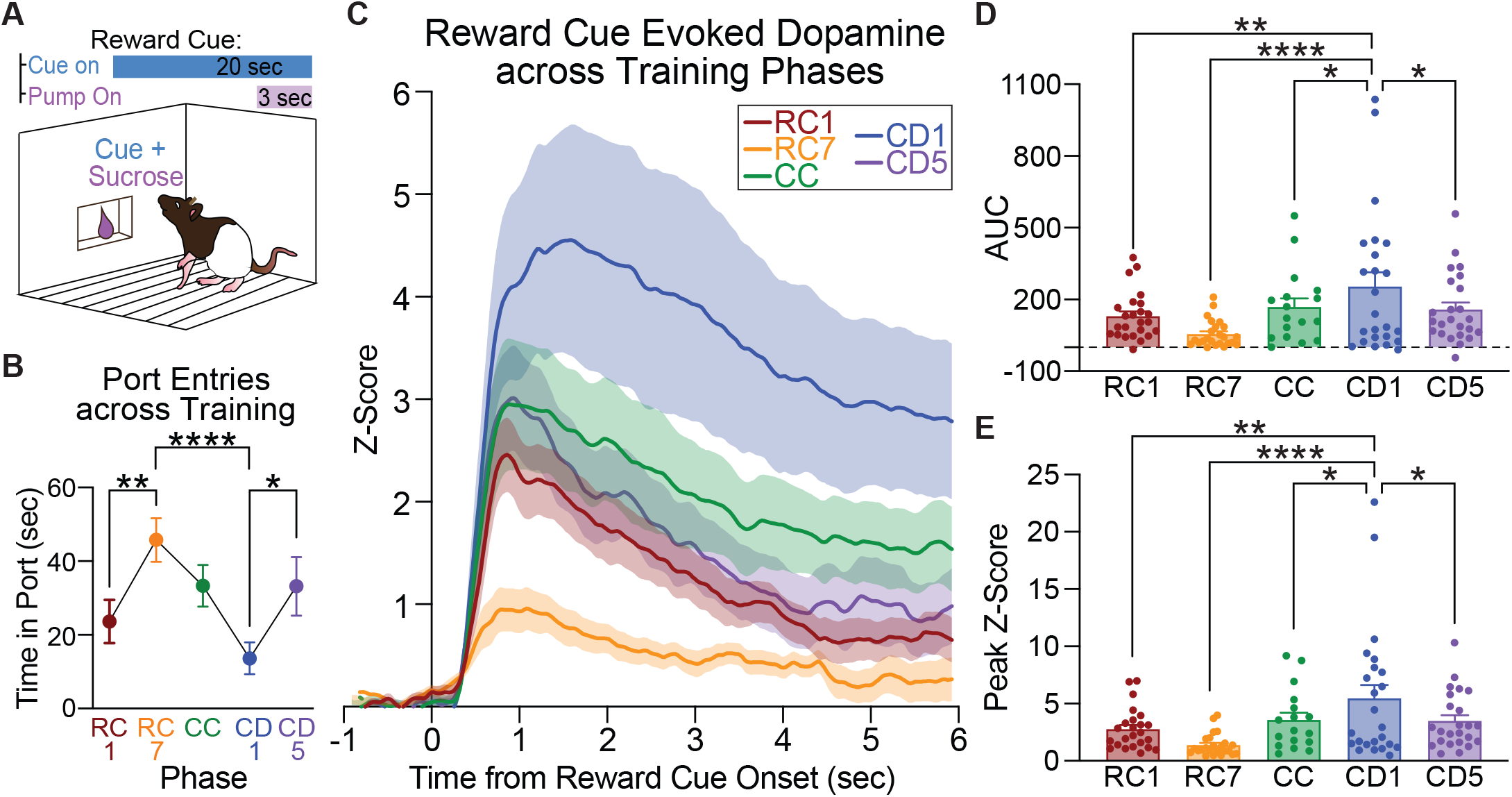
Reward-cue evoked dopamine signals dynamically rescale with changing emotional uncertainty. A) Throughout all of the multi-phase Pavlovian cue discrimination paradigm, rats learned a 20-s auditory cue predicted a 3-s delivery of sucrose into the reward port. B) In all phases, rats spent time in the reward port during reward cue presentations. Port entry behaviors increased during reward conditioning and cue discrimination, but generally transiently decreased as new cue types were introduced. C) Here, the reward cue evoked BLA dopamine responses across the experiment phases are plotted together. These signals consisted of a mild to moderate peak with varying levels of sustained activity. F) AUC comparisons of the first 2-s after reward cue onset and E) peak measures of the BLA dopamine signals show the reward cue elicited the highest amount of dopamine during the first day of cue discrimination compared to all other phases. RC = reward conditioning. CC = compound conditioning. CD = cue discrimination. Data represent mean ± SEM. *p<0.05, **p<0.01, ****p<0.0001.

During the first discrimination session, the reward cue elicited a more sustained signal (**Fig 5D**) and a higher peak value (**Fig 5E**) compared to the reward cue signals on reward conditioning day 1 (AUC: p=0.0034; peak value: p=0.0014), reward conditioning day 7 (AUC: p<0.0001; peak value: p<0.0001), compound conditioning (AUC: p=0.0167; peak value: p=0.0157), and the final discrimination (AUC: p=0.0408; peak value: p=0.0357). Further, on the last discrimination day, the reward cue signal was greater than the reward conditioning day 7 signal in terms of AUC (p=0.0234) and peak value (p=0.0215). These data support the notion that BLA dopamine scales with emotional uncertainty, in this case when new cues need to be disambiguated. The first day of the cue discrimination phase had the highest reward cue induced dopamine response (**Fig 5C**), but also had the lowest port entry behaviors (**Fig 5B**). Moreover, while time in the reward port increased across the cue discrimination phase (**Fig 5B**), the reward cue traces diminished (**Fig 5C**). These data reinforce the notion from our correlation data that BLA dopamine response does not encode behavioral expression, instead dynamically reflecting the relative emotional intensity of sensory cues.

### Bias in extinction of reward versus fear cue-evoked BLA dopamine dynamics

We next conducted extinction of cue discrimination (**Fig 6A**), wherein the reward and fear cues were presented without sucrose or footshock (**Fig 6B**). Conditioned behavioral responses extinguished quickly. When compared to the final day of cue discrimination, time spent in the reward port during reward cue presentations significantly decreased (**Fig 6C**; (t(19)=4.134, p=0.0006). Within extinction training, the time spent in the reward port during the reward cue presentations was generally greater than the fear cue presentations (**Fig 6C**; F(1,46)=8.674, p=0.0050), but this difference was gone by the third extinction session (**Fig 6C**; p=0.2148). Concurrently, cue-induced freezing behavior to the fear cue was higher than the reward cue (**Fig 6D**; F(1,46)=18.16, p<0.0001) throughout all of extinction training, but this cued freezing behavior discrimination decreased across sessions (**Fig 6D**; session x cue type interaction F(2,78)=5.882, p=0.0042). This effect was especially blunted by the end of extinction as freezing to the fear cue was significantly lower on day 3 compared to day 1 (p=0.0029). Females and males had similar cue-induced freezing during extinction training, however, different cue types prompted varying levels of freezing between the sexes, as reward cue induced freezing decreased in females, but was maintained in males during extinction (session x sex interaction, F(2,37)=4.852, p=0.0135).

**Fig 6.**
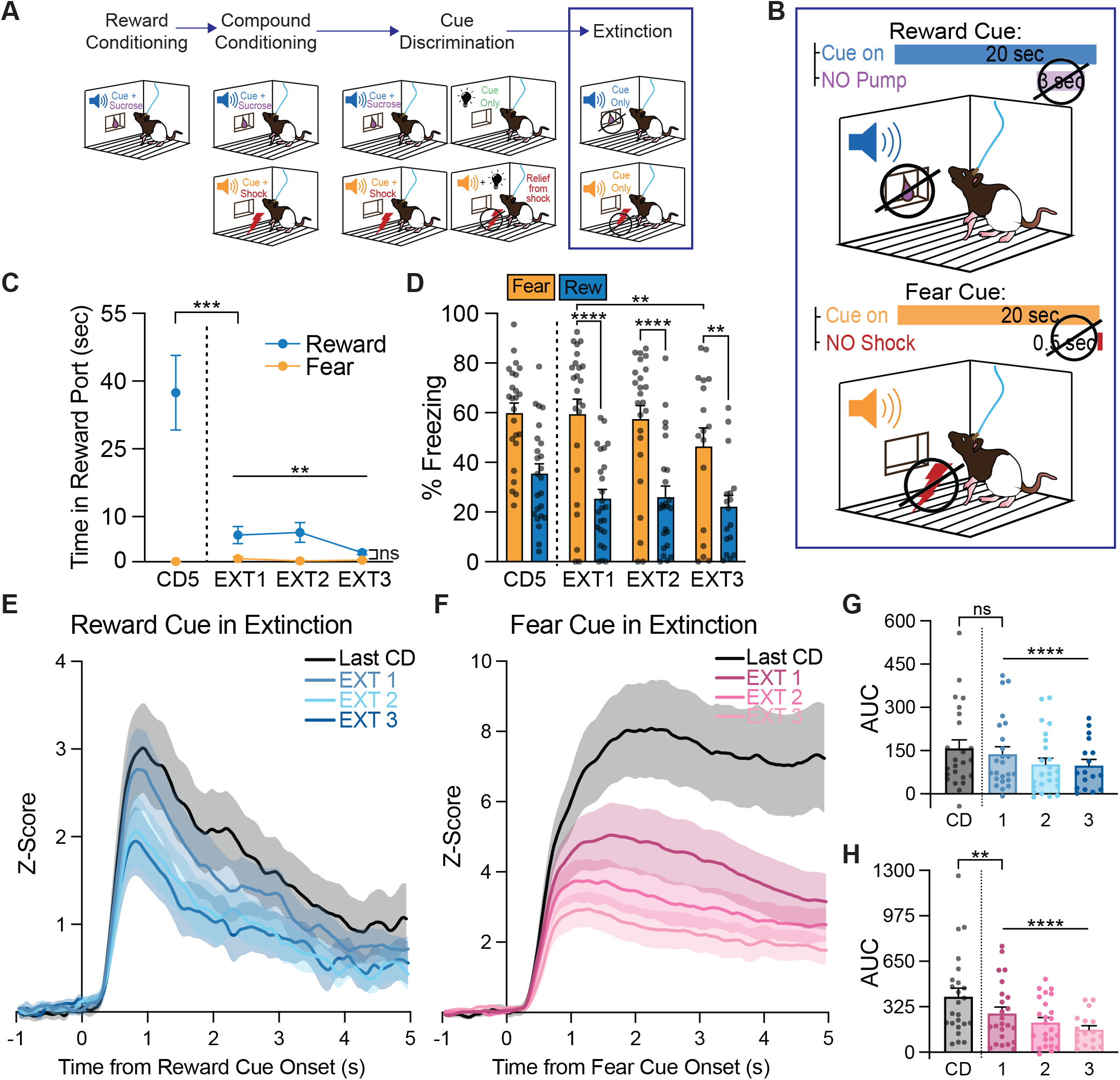
Bias in extinction of reward versus fear cue-evoked BLA dopamine dynamics. A) Schematic of the multi-phase Pavlovian cue discrimination paradigm the rats were trained on (n=24; 11F, 13M). Behavior and photometry data from the extinction training phase are shown. B) The reward cue and fear cue were presented without their previously paired outcomes (sucrose or footshook). C) Compared to the last day of cue discrimination, port entry behaviors diminished during the reward cue when sucrose is omitted and remained absent during fear cue presentations. D) Cue induced freezing behavior decreased for the fear cue across extinction training, while freezing responses remained low for the reward cue. E) The reward cue continued to elicit moderate BLA dopamine dynamics during the first extinction with similar levels to the final discrimination session, but began to lessen throughout extinction training. F) The fear cue dopamine trace immediately decreased from the levels seen in the discrimination phase, and continued to temper down until the final extinction training day. G) Within extinction, AUC values decreased across extinction for the reward cue, H) and also decreased for the fear cue. This pattern was also seen when comparing the fear cue AUC values from extinction day 1 to the final cue discrimination session. Data represent mean ± SEM. *p<0.05, **p<0.01, ***p<0.001, ****p<0.0001. CD=cue discrimination.

Behavioral changes in cued locomotion levels were similar in both females and males (**Fig S2**). However, while the qualitative signal patterns and dynamics of the cue-evoked BLA dopamine established during discrimination training were partially maintained throughout all three extinction sessions for the reward cue (**Fig 6E**) and fear cue (**Fig 6F**), there were significant changes in the magnitude of evoked signals and encoding of both cue types. Dopamine responses decreased in response to the reward cue throughout extinction training (**Fig 6E,G**; F(2,39)=12.35, p<0.0001; day 1 vs day 2, p=0.0139 and day 3, p<0.0001), but not when comparing the first extinction session to the final discrimination session (t(23)=1.403, p=0.1740). Alternatively, the fear cue-evoked signal quickly decreased relative to cue discrimination (t(23)=3.125, p=0.0048) and continued to decrease as extinction training progressed (**Fig 6F,H**; F(2,39)=19.28, p<0.0001; day 1 vs day 2, p=0.0187 and day 3, p<0.0001).

### BLA dopamine scales with threat intensity but not fear expression

Given that we saw larger BLA dopamine signals to fear-related cues during compound conditioning (**Fig 3H**) and cue discrimination (**Fig 4E,F**), we sought to understand if BLA dopamine would also scale when rats were tasked with discriminating cues associated with different levels of aversive valence. During a fear scale discrimination behavioral paradigm, rats were presented with intermingled fear cues within the same session (**Fig 7A**): a low shock cue (20-s auditory tone paired with a 0.2mA, 0.5s footshock) and a higher, moderate shock cue (20-s auditory tone paired with a 0.4mA, 0.5s footshock), presented 5 times each per session. Rats froze to both cues, and this freezing grew across sessions (**Fig 7B**; F(1,9)=44.91, p<0.0001). By the final day of fear scale discrimination rats exhibited higher freezing levels compared to first day levels for the high shock cue (p=0.0014) as well as the low shock cue (p=0.0164). However, freezing was not significantly modulated by shock cue type (F(1,9)=0.09096, p=0.7698).

**Fig 7.**
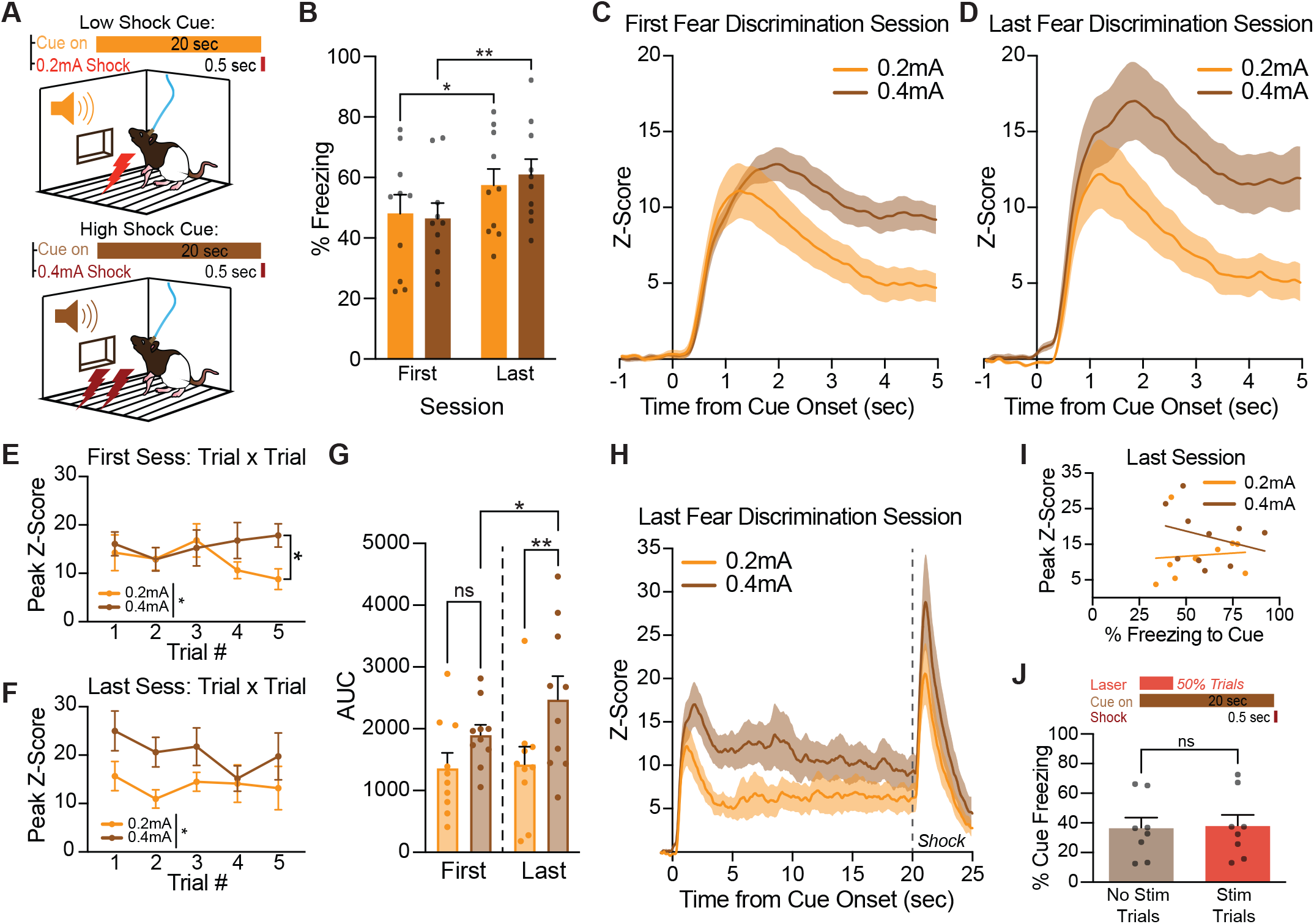
BLA dopamine tracks threat intensity but not fear expression. A) Rats (n=10; 4F, 6M) underwent scaling fear discrimination, where two 20-s auditory cues were presented intermingled in each training session, one cue culminated with a 0.2mA footshock (“Low Shock Cue”) and the other cue with a 0.4mA footshock (“High Shock Cue”). B) Cue-induced freezing behavior was high for both fear cue types. C) Upon the first discrimination sessions, both fear cues evoked robust BLA dopamine responses that were sustained in nature. D) By the last fear discrimination session, both fear cues maintained their robust and sustained BLA dopamine responses. E) There were five cue trials for each fear cue type. Within the first session, peak values of the dopamine traces began to diverge in magnitude, such that by the final cue presentation, the peak of dopamine trace for the high shock cue was larger than the peak of the low shock cue. F) By the last fear discrimination session, both fear cues maintained their robust and sustained BLA dopamine responses. G) Within the last discrimination session, the high shock cue presentations elicited higher dopamine dynamic peak values compared to the low shock cue presentations. H) BLA dopamine traces for the full 20-s cue presentations of the low and high shock cues. The high shock cue remains elevated compared to the low shock cue throughout the entire cue presentation as well as during the brief shock at 20-s. I) On the final fear scale discrimination session, neither the low shock cue dopamine signals (orange) nor the high shock cue dopamine signals (brown) were correlated with cue-induced freezing behaviors. J) A separate group of TH-cre rats (n=8; 3F, 5M) underwent fear conditioning paired with optogenetics. Dopamine terminals in the BLA were excited (5-sec, 20 Hz) at the onset 50% of fear cue presentations, which did not affect expression of freezing behavior. Data represent mean ± SEM. *p<0.05, **p<0.01.

Both fear cues evoked large and sustained BLA dopamine signals during the first fear discrimination session (**Fig 7C**). When comparing the average cue peak dopamine response for each trial, we saw a rapid distinction between the higher shock and lower shock cues emerge by the end of the first session, where the final high shock cue was significantly larger than the final low shock cue (**Fig 2E**; t(8)=3.126, p=0.0141).

The robust and sustained BLA dopamine responses were preserved during the last fear discrimination session for both the low and high fear cues (**Fig 7D**). The high shock versus low shock distinction persisted and was clearer across the last fear discrimination session. The high shock cue evoked higher dopamine peak values compared to the low shock cue presentations across trials in the final session (**Fig 7F**; F(1,9)=9.216, p=0.0141). Overall across training sessions the high shock cue dopamine AUC was greater than the low shock cue AUC (**Fig 7G**; F(1,9)=11.15, p=0.0087). This was driven by a trend for a larger high shock cue AUC in the first discrimination session (p=0.0637), which was significantly larger than the low shock cue by the final session (p=0.0483), and the high shock cue signal on the first session (p=0.0025). The low shock cue AUC did not significantly change between the first and last fear discrimination sessions (p=0.8121). Looking at the entire cue window, we saw both shock cues elicited prolonged BLA dopamine signals that persisted throughout the entire 20-s cue presentations. The high shock cue remained elevated in comparison to the low shock cue during the cue period as well as the footshock (**Fig 7H**). Despite this, cue evoked dopamine signals were not significantly correlated with freezing behavior for either the low or high shock cue (**Fig 7I,J**; Pearson correlation, low: p= 0.8169, high: p= 0.4391).

VTA dopamine neuron activation can facilitate fear learning (Jo et al., 2018) and specific inhibition of VTA-BLA dopamine neuron projections can disrupt fear memory recall (Tang et al., 2020). While we established fear cue presentations prompt strong BLA dopamine signals (**Fig 3I; Fig 4E,F; Fig 7C,D,H**), across studies, we did not find consistent correlations between fear cue evoked dopamine and conditioned freezing (**Fig 3L; Fig 4J; Fig 7I**). To further explore this, we tested the impact of exogenously increasing BLA dopamine levels, in TH-cre rats. We found that optogenetic excitation of VTA-BLA dopamine terminals during fear cue presentations did not affect freezing behavior (**Fig 7J**; t(7)=1.081, p=0.3156). Together with our above results, these data suggest that elevated cue-evoked dopamine reports the state of emotional intensity but does not acutely drive behavioral expression of emotional responses.

## DISCUSSION

We investigated how the BLA dopamine system is engaged as rats learn appetitive, aversive, and ambiguous associations. First, we show that VTA dopamine neuron stimulation engages robust BLA dopamine signaling that is not intrinsically reinforcing. Second, appetitive and aversive events, and their predictive cues, evoke BLA dopamine in simple cue conditioning paradigms, where signals scale with stimulus intensity but not reward value. Critically, cues evoked dopamine in a pattern that did not match the strength of learning, as signals were larger early and diminished across training, but rebounded when the emotional uncertainty of the sensory environment changed. As we increased the complexity of the learning context, where cues signaling threat, reward, safety, and neutral associations were intermingled, the level of cue-evoked BLA dopamine reported the relative emotional salience: fear and safety cues elicited larger dopamine signals than either reward or neutral cues. Finally, we show that BLA dopamine signals scale with the intensity of conditioned threats, but not the expression of fear. Together, our results show that BLA dopamine responds to salient cues, regardless of valence, and the magnitude and persistence of these signals dynamically adjusts according to the emotional relevance of the stimulus within a given learning context. These unique patterns highlight a dopamine system that encodes sensory state transitions in a way that is independent of associative strength or value. Our data help expand conceptual frameworks of emotional learning and offer a number of new insights into amygdala and dopamine functions (Tye and Janak, 2007; Gründemann and Lüthi, 2015; Sengupta et al., 2018; Zhang and Li, 2018; Corder et al., 2019; Gründemann et al., 2019; Zhang et al., 2021; Kong et al., 2024, 2025; Ottenheimer et al., 2024; Wojick et al., 2024).

### BLA dopamine signals do not predominantly reflect value or cue associative strength

We show that VTA dopamine neurons evoke strong dopamine release in the BLA (Brinley-Reed and McDonald, 1999; Takahashi et al., 2010; Lutas et al., 2019). While activation of VTA dopamine projections to the striatum is well documented to create associative learning and drive behavioral reinforcement (Witten et al., 2011; Kim et al., 2012; Steinberg et al., 2014; Wolff and Saunders, 2024), we found that VTA-BLA dopamine signaling is not reinforcing. Further, BLA dopamine levels increased during both aversive and appetitive events (Inglis and Moghaddam, 1999; Sias et al., 2024), and these signals scaled with stimulus intensity, but not value. Together, this suggests that dopamine release in the BLA does not directly drive learning, nor does it signal the positive or negative aspects of stimuli per se, but is instead involved broadly in the encoding sensory transitions.

Given that dopamine inputs to the BLA do not predominantly signal valence or value, the encoding of stimulus properties to represent valence instead is likely processed through different BLA efferent pathways to downstream regions and/or via different populations of BLA neurons (Stuber et al., 2011; Namburi et al., 2015). The BLA contains valence-sensitive clusters of neurons, where positive and negative conditioned stimuli are preferentially encoded (Paton et al., 2006; Gore et al., 2015; Namburi et al., 2015; Beyeler et al., 2016, 2018; O’Neill et al., 2018; Zhang and Li, 2018; Gründemann et al., 2019; Zhang et al., 2021). These ensembles are anatomically interspersed but are largely separate and similarly sized populations. While there is clear evidence that some BLA neurons have inherent bias for representation of appetitive or aversive stimuli, the differential encoding of valence information among BLA neurons is dynamic with learning (Rogan et al., 2005; Tye et al., 2008; Sang-ha et al., 2013). Plasticity driving this discrimination is supported by a complex array of circuit inputs and local communication within the amygdalar complex (Saddoris et al., 2005; Malvaez et al., 2015; Ottenheimer et al., 2024; Favila et al., 2025; Hinz et al., 2025). Dopamine input also likely plays a key role, as recent studies and computational models suggest (Hagihara et al., 2021; Lutas et al., 2022; Kuznetsov, 2025; Zhang et al., 2025). While additional experiments are needed to investigate this directly, we hypothesize that moment to moment, sensory-guided BLA dopamine fluctuations interface with distinct BLA neural ensembles for dynamic disambiguation of state.

Notably, across our experiments, BLA dopamine signal patterns did not fit clearly with associative learning or prediction error frameworks - cue-evoked dopamine responses were generally larger at the beginning of a conditioning phase, before strongest cued behavioral performance. Further, these signals were engaged independent of valence and value, as neutral, threat, reward, and safety cues all evoked a positive excitation of dopamine release. We also found no consistent correlations between cue-evoked BLA dopamine signals and behavioral expression. Overall, these patterns are in stark contrast to classic notions of the striatal dopamine system, where learning rate and prediction are reflected in incrementally accrued cue responses, and value is represented by negatively biased signals to aversive stimuli (Roitman et al., 2008; Badrinarayan et al., 2012; McCutcheon et al., 2012; Oleson et al., 2012; Lammel et al., 2014; Patriarchi et al., 2018; de Jong et al., 2019; van Elzelingen et al., 2022a; Bornhoft et al., 2025; Lopez and Lerner, 2025). This will require further study, but a lack of clear connection between BLA dopamine signals and behavioral expression in our data is perhaps consistent with studies showing that the BLA is not essential for expression of some forms of learning (Parkinson et al., 2000; Setlow et al., 2002; Izquierdo and Murray, 2007).

While there are other possible BLA dopamine input sources, such as from the substantia nigra, periaqueductal gray, and dorsal raphe nucleus (Pinard et al., 2008; Mingote et al., 2015; Poulin et al., 2018; Aksoy-Aksel et al., 2021; Zhao et al., 2022), we hypothesize that input from VTA dopamine neurons is responsible for signaling the cued state changes we see in the BLA. This notion is supported by previous studies. For example, lesions of VTA dopamine neurons impair BLA cue attentional signals (Esber et al., 2012). VTA to BLA dopaminergic terminals are activated during learning (Lutas et al., 2019) and they are necessary for some aspects of reward and fear learning (Fadok et al., 2009; Sias et al., 2024). Further, there is heterogeneity of stimulus response profiles within populations of VTA dopamine neurons (Cohen et al., 2012; Schultz, 2016; Morales and Margolis, 2017; Park and Moghaddam, 2017; Jo et al., 2018; Saunders et al., 2018; Cai et al., 2020; Elum et al., 2024) but a relatively anatomically uniform distribution of VTA to BLA projection neurons (Mingote et al., 2015; Tang et al., 2020). While it is unclear if they project to the BLA, a subset of VTA dopamine neurons are activated by both fear cues and non-predictive cues, suggesting the capacity to signal more general salience information (Jo et al., 2018). Collectively, our data reveal a unique dopamine signaling profile, which adds new insight to the growing understanding that dopamine neurons contribute heterogeneous information across projection sites (McCutcheon et al., 2012; Lammel et al., 2014; Kutlu et al., 2021; van Elzelingen et al., 2022a, 2022b; Bornhoft et al., 2025).

### BLA dopamine dynamically tracks cue salience and emotional ambiguity

Our findings suggest that rather than reporting the strength of learning, valence, value, or behavioral vigor, BLA dopamine transmits information related to the current emotional salience and ambiguity of sensory transitions, with signals biased towards the most motivationally important stimulus in the environment. We found that dopamine signals in response to reward cues dynamically shrunk or grew not as a function of behavioral expression as learning progressed but instead tracked the emotional uncertainty of the learning context. Reward cue-evoked dopamine release attenuated over simple reward conditioning sessions, but when additional cues were added, requiring discrimination and increasing the emotional complexity of the environment, reward cue signals rebounded above those previously low levels. This qualitative pattern, which is again in contrast to classic associative dopamine patterns in the striatum, underscores the notion that BLA dopamine signals salient shifts in state and local stimulus context, rather than simply denoting either stimulus novelty or accrued learning. Consistent with this, past studies using microdialysis show that tonic dopamine levels in the BLA slowly increase while rats undergo discrimination learning (Hori et al., 1993), and pharmacologically boosting BLA dopamine enhances the early acquisition of a stimulus-reward association (Hitchcott et al., 1997). Further, in humans, dopamine is released in the amygdala during shifts from rest to working memory (Fried et al., 2001), suggesting that this system is re-engaged when memory of past associations are needed (Sias et al., 2021, 2024). In our data, we also saw the clear change over time in BLA dopamine in response to the safety cue, where signals were initially larger and comparable to the fear cue, but decreased as discrimination learning progressed. These patterns support the notion that BLA dopamine is not purely biased towards negatively-valenced events but, more broadly, is invigorated and then reinvigorated when new stimuli in the environment are introduced. Collectively, our results and other important recent studies (Sias et al., 2024), support a framework where BLA dopamine contributes more directly to flexible state or model-based learning, rather than reward prediction error, incentive motivation, or reinforcement (Prévost et al., 2013; Saez et al., 2015).

We found that cues bearing greater emotional weight not only evoked larger peak BLA dopamine at cue onset, but also more sustained increases throughout the entire cue period. Fear cue-evoked signals were larger and broader, while reward and neutral cue signals were smaller and more transient. Interestingly, the safety cue evoked an intermediate response that was sustained but smaller than the fear cue signal. In line with this, when fear cues predicting different threat levels were intermixed, dopamine responses were more sustained to whichever threat was larger in the moment. This scaling of sustained dopamine matched the perceived emotional vigilance, a pattern that is again distinct from cue-evoked dopamine signals in striatum, which are typically composed of rapid phasic fluctuations (Roitman et al., 2008; Badrinarayan et al., 2012; McCutcheon et al., 2012; Oleson et al., 2012; Lammel et al., 2014; Patriarchi et al., 2018; de Jong et al., 2019; van Elzelingen et al., 2022a; Bornhoft et al., 2025; Lopez and Lerner, 2025). Mechanistically, sustained BLA dopamine patterns may be supported by the relatively low levels of the dopamine transporter (DAT) and slower dopamine clearance rates within the amygdala, compared to the striatum (Revay et al., 1996; Holleran et al., 2020). Dopamine signaling in the BLA is known to affect long-term potentiation, and engenders complex effects on local and projection neuron populations, partially via modulation of glutamatergic inputs (Rosenkranz and Grace, 1999; Bissière et al., 2003). Dopamine neurons projecting to the BLA exhibit unique physiological characteristics (Ford et al., 2006; Margolis et al., 2006), preferentially targeting BLA projection neurons containing the D1-type dopamine receptors (D1DR) (Scibilia et al., 1992; Kröner et al., 2005; Muller et al., 2009; Takahashi et al., 2010; Aksoy-Aksel et al., 2021). Optogenetic activation of dopamine release increases cyclic AMP activity in BLA D1DR neurons and enhances their response to sensory inputs (Lutas et al., 2022), while pharmacological activation of BLA D1DR signaling facilitates contextual fear learning (Heath et al., 2015), freezing behavior (Guarraci et al., 1999), reward seeking and learning (Tye et al., 2010), as well as safety cue-induced fear suppression (Ng et al., 2018). Our results motivate future studies aimed at dissecting the contribution of stimulus-evoked BLA dopamine sig-nals of different duration and intensity on plasticity in the BLA. One prediction, motivated by our data, is that sustained levels of dopamine during discrimination of different cues may support stimulus context encoding on longer timescales. Given that BLA dopamine levels are dynamically modulated by changes in affective and homeostatic state, these signals could contribute to scaling of value and memory representations among BLA neurons (Maren, 2005; Grove et al., 2022; Hinz et al., 2025; Lesuis et al., 2025).

Here, we showed BLA dopamine release tracks reward, fear, discrimination, and extinction learning. Taking the classic view that the BLA processes sensory stimuli on both the proximal and environmental levels to support behavioral outputs, our results imply that BLA dopamine signals are involved in valence discrimination via a transmission bias that reports the relative emotional saliency and uncertainty of the moment. These signals are dynamically engaged in learning contexts where relative stimulus importance must be rapidly disambiguated. Our results support a framework where the BLA dopamine system, in contrast to classic striatal dopamine frameworks, serves as a general, sensory state-guided saliency signal, to support emotional learning.

## Funding and Acknowledgements

The authors have no biomedical financial interest or potential conflicts of interest. This work was supported by NIH grants F31 DA060004 (MAB), P30 DA048742 (MJT), and R00 DA042895, R01 MH129370, R01 MH129320, and R01 DA057292 (BTS). We thank all members of the Thomas and Saunders labs for technical support and feedback.

## METHODS

### Subjects

Male and female Long Evans rats (n=41; 19F, 22M) were used. All rats weighed 250-500g at the time of surgery and were 4-9 months old at the time of experimentation. Experimental procedures were approved by the Institutional Animal Care and Use Committee at the University of Minnesota and were carried out in accordance with the guidelines on animal care and use of the National Institutes of Health of the United States.

### Surgical Procedures

Rats were anesthetized with 5% isoflurane and placed in a stereotaxic frame, after which anesthesia was maintained at 1-3%. Saline, carprofen anesthetic (5 mg/ kg), and cefazolin antibiotic (70 mg/kg) were administered subcutaneously prior to start of surgery. The top of the skull was exposed and holes were made for viral infusion needles, optic fiber implants, and 4 skull screws. A total of 0.5 µL of a virus coding for the dopamine biosensor dLight 1.3b (AAV5-hSyn-dLight1.3b, University of Minnesota Viral Vector and Cloning Core) was infused unilaterally into the BLA (2.7 AP, +/-4.9 ML, -8.2 DV). In a separate cohort of rats, a total of 1.0 μL of virus (pAAV5-Syn-FLEX-Chrimson-tdTomato, Addgene) was infused unilaterally into the VTA (−5.6 AP, +/− 0.7 ML, −8.0 DV), while dLight1.3b injections were targeted to the ipsilateral BLA. In another cohort of rats, a total of 1.0 μL of virus (pAAV5-EF1a-double floxed-hChR2(H134R)-EYFP-WPRE-HGHpA, Addgene) was infused unilaterally into the VTA. Each virus was delivered at 0.1µL/min. Immediately after injection, the needle was raised 100 µm and then left in place for an additional 10 min to allow for diffusion. Optical fibers (9mm length, 400µm diameter, Doric Lenses) for photometry recordings were chronically implanted into the BLA (2.7 AP, +/-4.9 ML, -7.9 DV) in the same hemisphere as the viral infusion. For optogenetic stimulation, optical fibers (10mm length, 300μm diameter, Thorlabs) were implanted above the VTA (−5.6 AP, +/− 0.7 ML, −7.9 DV) and/or the BLA (2.7 AP, +/-4.9 ML, -8.0 DV) in the same hemisphere of viral injections. All coordinates are in mm relative to bregma and skull surface. Implants were secured to the skull with dental acrylic (Lang Dental) applied around the skull screws and the base of the ferrules containing the optic fibers. At the end of all surgeries, topical anesthetic and antibiotic ointment were applied to the surgical site, rats were placed on a heating pad and monitored until they were ambulatory. After surgery rats were individually housed with ad libitum access to food and water on a 0800 to 2000 light/dark cycle (lights on at 0800), given carprofen (5 mg/kg) and cefazolin (70 mg/kg) for the first 3 days following surgery and weighed and monitored for 6 days. Photometry recordings began at least 3 weeks after surgery. Optogenetic manipulations started at least 6 weeks after surgery.

### Fiber Photometry Recordings

Behavioral testing days were paired with fiber photometry recording to measure dopamine signals (n=32; 15F, 17M) in the BLA. The fiber photometry system was outfitted with optical components from Doric Lenses (Québec, Canada). This included fluorescence mini-cubes to transmit concurrent light streams from LEDs, 465nm and 405/415nm, that were sinusoidally modulated at 211Hz and 330Hz, respectively. The 465nm channel stimulated fluorescence from the dopamine-dependent dLight, while the 405/415nm served as an isosbestic reference that excites dLight in a dopamine-independent manner. Demodulating the brightness produced by both channels allows for reliable correction of movement or photobleaching artifacts. Before each photometry recording both LED currents were set to ∼50μW. Fluorescence was transmitted from the fiber tip over the BLA back to the mini-cube via an optic cable, where it was passed through a GFP emission filter, amplified and directed into a high sensitivity photoreceiver (Newport, Model 2151) for signal processing. Meanwhile, the session was controlled and data was acquired via Synapse software running on a Tucker-Davis Technologies (TDT, Alachua, FL) real-time processor (RZ5P or RZ10X), as reported in previous studies (Engel et al., 2024; Bornhoft et al., 2025). Synapse software on the processor calibrated the output of each LED channel and simultaneously recorded photometry signals, which were being sampled from the photodetector at 6.1 kHz.

To align the real-time neural signals to behavioral (e.g., port entries) and environmental (e.g., cue presentations) occurrences, TTL signals from the operant chambers (Med Associates) were sent to the RZ5P processor and timestamped for post-hoc analysis of the photometry data. Video recordings were also taken and integrated with the photometry and behavior data via the Synapse software.

### Optogenetic Stimulation and Behavior

#### Optogenetic stimulation

These studies used 589-nm lasers (OptoEngine and Dragon Lasers) and 473-nm lasers (OptoEngine). Light output included individual 5ms light pulses for either 5Hz or 20Hz stimulation frequency. Light intensity was adjusted to be approximately 2 mW/ mm2 at the tip of the intracranial fiber during individual pulses (∼10-mW constant power). The optic tethers connected to the rats were sheathed in a lightweight armored jacket to block visible light transmission and reinforce the cable to prevent breakage. Laser pulses were initiated from Med-PC TTL pulses and patterned via Synapse software that also controlled photometry and video data acquisition. Fiber photometry recordings were concurrently collected on every optogenetic training day.

#### Fiber Photometry and Optogenetics

To understand how VTA TH-neuron activity influences dopamine dynamics in the BLA, in vivo optogenetics was paired with fiber photometry in TH-Cre rats (n=8; 4F, 4M). Rats were first acclimated to the behavioral chambers (Med Associates) and optic cable tethering in a ∼30-min habituation session. During VTA TH-cell body stimulation, an optic tether was attached over the VTA implant to deliver optogenetic stimulation while another was attached to the BLA implant for dLight fiber photometry data collections. In separate sessions, rats were tethered to the optic cable and presented with 8-10 laser stimulations on a 120-s fixed time schedule. The stimulation parameters were (1) 20Hz stimulation for 2 s, (2) 5Hz stimulation for 5 s, and (3) 20Hz stimulation for 5 s. These stimulation presentations were un-cued and not paired with any external stimulus. The same stimulation parameters and sessions were repeated for VTA TH-neuron terminal stimulation over the BLA. Here, the optic tether connected to the BLA implant concurrently delivered the laser stimulation, while collecting dLight fiber photometry fluorescence signals.

#### Intracranial self-stimulation (ICSS)

We expressed Chrimson in VTA dopamine neurons in TH-Cre rats (n=14; 6F, 8M) and inserted an optic fiber in the BLA. Rats were placed in the testing chamber (Med Associates) and connected to an optic cable tether to receive optogenetic stimulation to target VTA dopamine terminals in the BLA. Two nose poke ports were positioned on one side of the chamber. Nose pokes in the active port resulted in a 2-s laser train (20Hz, 5-ms pulses, fixed-ratio 1 schedule) that coincided with the delivery of a tri-color light illumination cue. Laser stimulation parameters were chosen to align with the results from the fiber photometry studies above where 2-s, 20Hz VTA-BLA dopamine terminal stimulation prompted reliable dopamine dynamics in the BLA.

#### Optogenetic stimulation during fear conditioning

To investigate whether exogenously increasing BLA dopamine levels during a fear cue would impact freezing behaviors, Chrimson was virally expressed in VTA dopamine neurons in TH-cre rats (n=8; 3F, 5M) and an optic fiber was implanted over the BLA. Rats then underwent two days of fear conditioning, where a 20-s auditory cue was paired with a moderate footshock (0.4 mA) delivered from 20-20.5 s. There were 14 cue-shock presentations each session with each cue trial separated by an intertrial interval that averaged 115s (range: 100s-130s). Half of cue trials were paired with a 5s, 20Hz laser delivery concurrent with the fear cue onset to prompt VTA-BLA dopamine terminal activation. The stimulation trials were pseudorandomly delivered interwoven with the non-stimulation trials. Each session lasted ∼30 min.

### Pavlovian Behavioral Assays

All Pavlovian behavior training began at least 3-4 weeks after surgery. Med Associates chambers were outfitted with a syringe pump to deliver liquid sucrose reward, a magazine port equipped with an infrared beam to sense port entries, floor bars for the delivery of electrical shocks, speakers for the delivery of tone (high: 4500 Hz, low: 2900 Hz), and house lights for illumination. Cameras were positioned above each chamber to record movement throughout each session for post-hoc behavioral analysis. Port entry instances and durations were recorded by Med Associates software throughout the sessions. Chambers were cleaned with 70% ethanol and bedding was replaced after each session.

#### Cue Discrimination Paradigm

This Pavlovian cue discrimination task (n= 24; 11F, 13M) was adapted from previous studies (**Fig 4A**) (Sangha et al., 2013). Habituation: Training began with acclimation to the behavioral chambers and optic cable tethering in two ∼30-minute habituation sessions. During these sessions, rats were tethered to the fiber photometry cable and allowed to freely move and explore around the chamber.

#### Reward Conditioning

For the 7 following sessions, rats were presented with 25 auditory cue presentations lasting 20 s each on a 110s VT schedule, (range: 90-130 s). The auditory cue was either a pulsing low tone (repeating cycle of 0.2 s on and 0.2 s off) or a continuous high tone counterbalanced across rats. Each reward cue presentation was paired with a liquid reward (10% sucrose) delivered pseudorandomly between 10 s and 20 s after the reward cue onset. Each session lasted ∼55 min.

#### Cue Habituation

To acclimate the rats to the remaining cues, during an 8th reward conditioning session (25 auditory reward cue+sucrose presentations), the rats were pseudorandomly presented with noncontingent cues (5 house lights illuminations, 5 auditory (counterbalanced high or low) tones) lasting 20 s each. These cue presentations were not associated with any outcome. This session lasted ∼75 min.

#### Compound Conditioning

The next training session introduced the fear cue: 5 presentations of a 20-s auditory cue paired with a mild footshock (0.4 mA) delivered from 20-20.5 s. The fear cue presentations were pseudorandomly delivered intermixed with 25 reward cue presentations. This session lasted ∼65 min.

#### Cue Discrimination

After initial learning, the rats underwent the cue discrimination phase of the paradigm. Here, the rats had to distinguish between 4 different cues predicting various outcomes: the previously introduced reward cue (paired with sucrose) and (2) fear cue (paired with a footshock), as well as a (3) neutral cue with no paired outcome and a (4) compound cue (fear+neutral cue played concurrently) to signal a relief from shock. The high and low tones, as well as pulsatile versus continuous tones, were counterbalanced across rats for the reward and fear cues. House light illumination was always used for the neutral cue. The compound cue signaling relief from shock was always the fear cue (counterbalanced auditory cue) played concurrently with the neutral cue (visual cue) for 20 s. There were 45 total cue presentations in each training session (15 reward cues, 5 fear cues, 10 neutral cues, and 15 compound cues). Each cue lasted 20 s and was delivered in pseudorandom orders on a 110 s average VT schedule (range 90-130 s). Each animal had 4-6 cue discrimination sessions, each lasting ∼100 min.

#### Extinction

Behavioral testing concluded with 2-3 extinction sessions. Here, only the reward and fear cues were presented (20 presentations each; pseudorandom order) and both the reward and shock deliveries were omitted. These sessions lasted ∼85 min each.

#### Scaling Valenced Stimuli Presentations

##### Reward Scaling

Rats (n=8; 3F, 5M) were placed in Med Associates chambers while tethered to a fiber photometry cable. In one session, a 10% sucrose liquid reward was delivered via a syringe pump into the magazine port on the right side of the chamber. In a separate session, an Ensure liquid reward was delivered via a syringe pump into the magazine port on the left side of the chamber. For both reward types, there were 20 uncued, unsignaled reward presentations into the respective magazine port, which rats could enter at any time to consume the reward throughout the session. Both the left and right magazines were open during both reward sessions. Reward pumps were on for 3s and delivery occurred on a 105 s average VT schedule, (range: 90-120 s). Each session lasted ∼38 min.

##### Footshock Scaling

Rats (n=10; 3F, 7M) were placed in Med Associates chambers while tethered to a fiber photometry cable. Footshocks were delivered via electric currents through the floorbars. In one session, 0.2mA uncued, unsignaled 0.5 s footshocks were delivered. In a separate session, 0.4mA uncued, unsignaled 0.5-s footshocks were delivered. Sessions were conducted in a counterbalanced manner. In both sessions, there were 6 footshock presentations that were delivered on a 210 s average VT schedule, (range: 180-240 s). Each session lasted ∼18 min.

#### Fear Scaling Discrimination Paradigm

##### Single stimulus Fear Conditioning

Rats (n=10, 4F, 6M) underwent 2 distinct pavlovian fear conditioning sessions. One session had 5, 20 s auditory cue presentations paired with a lower footshock (0.2 mA, 0.5s) delivered at the end of the cue. A separate session had 5, 20 s auditory cue presentations paired with a higher footshock (0.4 mA, 0.5s) delivered at the end of the cue. The tones used were counterbalanced across rats. In one group, a continuous low tone was used for the 0.2mA cue, while a pulsing low tone (repeating cycle of 0.2 s on and 0.2 s off) was used for the 0.4mA cue. In the other group, a pulsing high tone (repeating cycle of 0.2 s on and 0.2 s off) was used for the 0.2mA cue, while a continuous high tone was used for the 0.4mA cue. In both sessions, the cues were delivered in pseudorandom order on a 110 s average VT schedule (range 90-130 s) and each session lasted ∼12 min.

##### Fear Discrimination

After initial learning, the rats underwent the cue discrimination phase of the paradigm. Here, the rats had to distinguish between the 2 different cues predicting the footshock outcomes: the previously introduced (1) low fear cue (paired with a 0.2mA) and(2)high fear cue (paired with a 0.4mA footshock). There were 10 total cue presentations in each training session (5 low fear cues and 5 high fear cues). Each cue lasted 20 s and was delivered in a pseudorandom order on a 110 s average VT schedule (range 90-130 s). Each animal had 3 fear discrimination sessions, each lasting ∼22 min.

### DeepLabCut-based pose tracking

Markerless tracking of animal body parts was conducted using the DeepLabCut Toolbox (DLC version 2.2.3 or 3.3.0) (Mathis et al., 2018) and analysis of movement features based on these tracked coordinates was conducted in Matlab R2024b (Mathworks). Videos (944 × 480 resolution) were recorded with a sampling frequency of 10-20 frames per second using TDT Synapse software with overhead cameras (Vanxse CCTV 960H 1000TVL HD Mini Spy Security Camera 2.8-12 mm Varifocal Lens Indoor Surveillance Camera).

#### DeepLabCut Model

Videos from this experiment were analyzed through a refined DLC network as described previously (Engel et al., 2024; Wolff and Saunders, 2024; Bornhoft et al., 2025). To create the network, the body parts labeled included the nose, eyes, ears, fiber optic implant, shoulders, tail base, and an additional three points along the spine. Features of the environment were also labeled, including the 4 corners of the apparatus floor and the two magazine ports. Briefly, 2090 frames from 35 videos (32 different animals, 3 experiments) were labeled and 807 outlier frames were relabelled to refine the network for the current study. Labeled frames were split into a training set (95% of frames) and a test set (5% of frames). A ResNet-50 based neural network was used for 1,030,000 training iterations. After final refinement, we used a p-cutoff of 0.85 resulting in training set error of 2.99 pixels and test set error of 3.68 pixels. This model was then used to analyze videos from the rats (n=26; 13F, 13M) who underwent the cue discrimination behavioral assay and the fear scale discrimination behavioral assay for all training sessions.

## Data Analysis

### Behavioral Analysis

Port entries, locomotion bouts, and freezing bouts were the primary behavioral output measures. Port entry frequency and duration was analyzed from MedPC software outputs and the generated TTL signals from Synapse. Video Scoring: Freezing data was first analyzed with hand-scoring of recorded videos from the cue discrimination training sessions. For each animal, freezing duration for each cue presentation was calculated during at least the first and last cue discrimination training session. The experimenter was blinded for both rat and session information. Cue-evoked freezing was timed for each cue three times and the average was calculated. DeepLabCut Analysis: Freezing bouts and locomotion bouts were also calculated from positional information tracked using DLC. Bodypart coordinates and confidence values for each video frame were determined by the DLC network and were analyzed with Matlab. Broadly, any frames with body part coordinates where confidence values were below 0.7 were excluded from further data processing. The average coordinates of fixed features of the operant chamber were used for analysis. A pixel-to-cm ratio was calculated for each processed video in order to transform pixel distances into physical chamber dimensions for subsequent distance traveled and speed analyses. The pixel distance of all 4 perimeter sides and both diagonal cross-sections were determined, divided by the actual distance in cm, then that quotient value is averaged to give the pixel-to-cm conversion for each video. Movement speed was calculated from the coordinates of the mid-back, for each video frame, the formula [distance moved (pix per cm) * framerate] was used to provide movement speed in cm/s. For analyzing data across recordings with different sampling rates, missing data points for the lower sampling rate sessions were filled in using linear interpolation with the interpol1 function (MATLAB). Interpolated data points were not used to calculate peak or mean speeds for analysis.

Freezing and locomotion detection was uniquely computed for each animal using a movement threshold determined by animal body size (distance between shoulder and lower back) and the predetermined optimal threshold for movement detection based on our operant chamber configuration (Collins et al., 2023; Engel et al., 2024; Wolff and Saunders, 2024; Bornhoft et al., 2025). This threshold value was used to detect fine movement in the face and head; for movement detection in the body, the limit was multiplied by 2 to account for larger body parts. Freezing bouts were defined as all visible body parts being below their corresponding movement threshold for any single video frame, while the transitions in and out of a movement bout were demarcated by a sliding window when the speed of 2 or more body parts exceeded the detection threshold for 0.3s. For all bouts, video frames where less than 2 body parts were visible were excluded. For freezing, bouts lasting under 1 s were excluded from bout analysis. Locomotion was detected using the speed of the implant and 4 locations on the spine. Locomotion bouts were determined if all visible body parts were above their corresponding movement threshold for any single video frame, while the transitions in and out of a movement bout were demarcated by a sliding window when the speed of 2 or more body parts exceeded the detection threshold for 0.3 s. For locomotion, bouts lasting less than 0.5s were excluded from bout analysis. For a subset of videos the filtered predictions were used and the average frame x frame movement across the fixed features of the environment was subtracted to remove jitter caused by camera movement. This restored freezing detection to similar levels as videos where camera movement was not a factor.

### Fiber Photometry Analysis

Photometry data were analyzed using a custom MATLAB pipeline and GraphPad Prism. First, the photometry signals (both 405nm/415nm isosbestic and 465nm) were low pass filtered and downsampled to 40Hz. The isosbestic signal was then aligned to the 465-nm signal by applying a least-squares linear fit and this fitted isosbestic signal was used to normalize the 465-nm signal, where ΔF/F = (465 nm signal – fitted 415-nm signal)/(fitted 415-nm signal). Timestamps for each video recording (recorded at 10-20 FPS) were recorded in the Synapse photometry data file. Cue presentations, port entries, sucrose pumps, and footshocks were assigned a TTL signal from Med-PC that was used to timestamp the photometry data file in the Synapse software. The normalized ΔF/F signals were extracted 5 s before and 25 s following each cue presentation and Z-scored to the 5 s pre-cue period for each trial to minimize the effects of drift in the signal across experiment duration. Normalized signals for reward consumption were extracted in the window from 2 s preceding to 4 s following each port entry and Z-scored to the 2 s pre-port entry. Port entry occurrences were binned into rewarded and unrewarded port entries. The first port entry after reward delivery with a duration of >0.5 s was designated a rewarded port entry. Normalized signals for shock responses were extracted in the window from 2 s preceding to 4 s following each shock delivery and Z-scored to the 2 s pre-shock delivery. Cue responses were detected in the average waveform for each animal in the 5 s window beginning at cue onset, while reward and shock responses used a 2 s window beginning at event onset. The maximum and minimum responses in each window were detected, and latency to the maximum or minimum was calculated relative to the start of the detection window. Area under the curve (AUC) values were calculated from the Z-scored traces by numerical integration via the trapezoidal method using the trapz function (Matlab).

### Statistics

Behavior and photometry signal data were analyzed with a combination of ANOVA models (one-way, two-way, and mixed effects), t-tests, and Pearson correlations. Post-hoc comparisons and planned t-tests were used to clarify main effects and interactions, with a Bonferroni correction applied. Photometry traces represent average values of sites/subjects, not individual trials. The data are expressed as the mean ± s.e.m. For all tests, the statistical significance was set at p<0.05.

## Histology

Rats were deeply anesthetized via i.p. injections of Fatal-Plus (2 ml/kg; Paterson Veterinary) and were transcardially perfused with cold phosphate buffered saline (PBS) and subsequent 4% paraformaldehyde (PFA). Brains were extracted and continued to be fixed in 4% PFA overnight (∼24 hrs) then were transferred to a cryoprotectant solution of 30% sucrose in PBS for at least 48 hrs. The tissue was cut into 50-μm sections on a freezing microtome (Leica Biosystems) and brain sections containing the BLA were mounted on microscope slides and coverslipped with Vectashield containing DAPI counterstain to validate viral expression and proper optic fiber placements. Images of the microscope slides were taken on a fluorescent microscope (Keyence BZ-X710) with a 4x and a 10x air immersion objective. Virus spreads and fiber positions were compared to the Rat Brain Atlas (Paxinos and Watson, 2007) and plotted with Adobe Illustrator.

### Immunohistochemistry

To visualize dopamine projections in the amygdala, we performed immunohistochemistry for tyrosine hydroxylase. Brain sections were washed in PBS and incubated with bovine serum albumin (BSA) and Triton X-100 (each 0.2%) for 20 min. 10% normal donkey serum (NDS) was added for a 30-min incubation, before primary antibody incubation (rabbit anti-TH, 1:500, Invitrogen) overnight at 4°C in PBS with BSA and Triton X-100 (each 0.2%). Sections were then washed and incubated with 2% NDS in PBS for 10 minutes and secondary antibodies were added (1:200 594 donkey anti-rabbit) for 2 hours at room temperature. Sections were washed 2 times in PBS and mounted with Vectashield containing DAPI.

## Acknowledgements

The authors have no biomedical financial interest or potential conflicts of interest. This work was supported by NIH grants F31 DA060004 (MAB), P30 DA048742 (MJT), and R00 DA042895, R01 MH129370, R01 MH129320, and R01 DA057292 (BTS). The authors thank all members of the Thomas and Saunders labs for technical support and feedback.

## Supplemental Material

**Fig S1.**
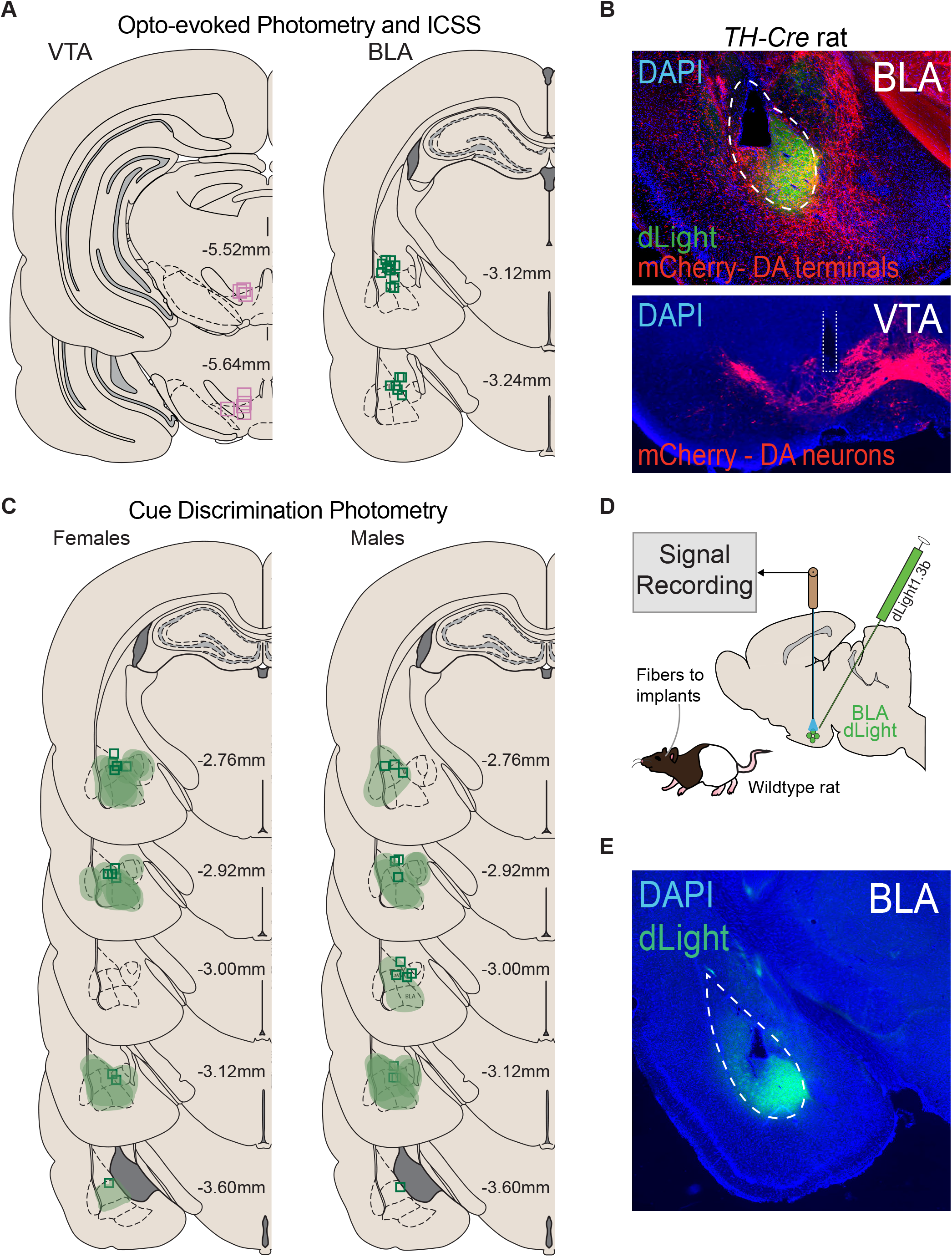
Histology for VTA-BLA optogenetic stimulation and dLight fiber photometry recordings. A) Optic fiber placements for rats used to measure optogenetically-evoked dopamine signals and ICSS behavior following optogenetic activation of VTA dopamine terminals in the BLA. B) Histology images showing BLA dopamine terminals (top) and VTA dopamine neuron cell bodies (bottom) expressing mCherry from a cre-dependent viral injection in the VTA of TH-cre rats. C) dLight 1.3b viral spread and optic fiber placements for the photometry cue discrimination experiments in female and male rats. D) Schematic of the fiber photometry approach with the dopamine sensor, dLight 1.3b, expressed in the BLA of freely moving rats. E) Representative image of dLight virus expression in the BLA.

**Fig S2.**
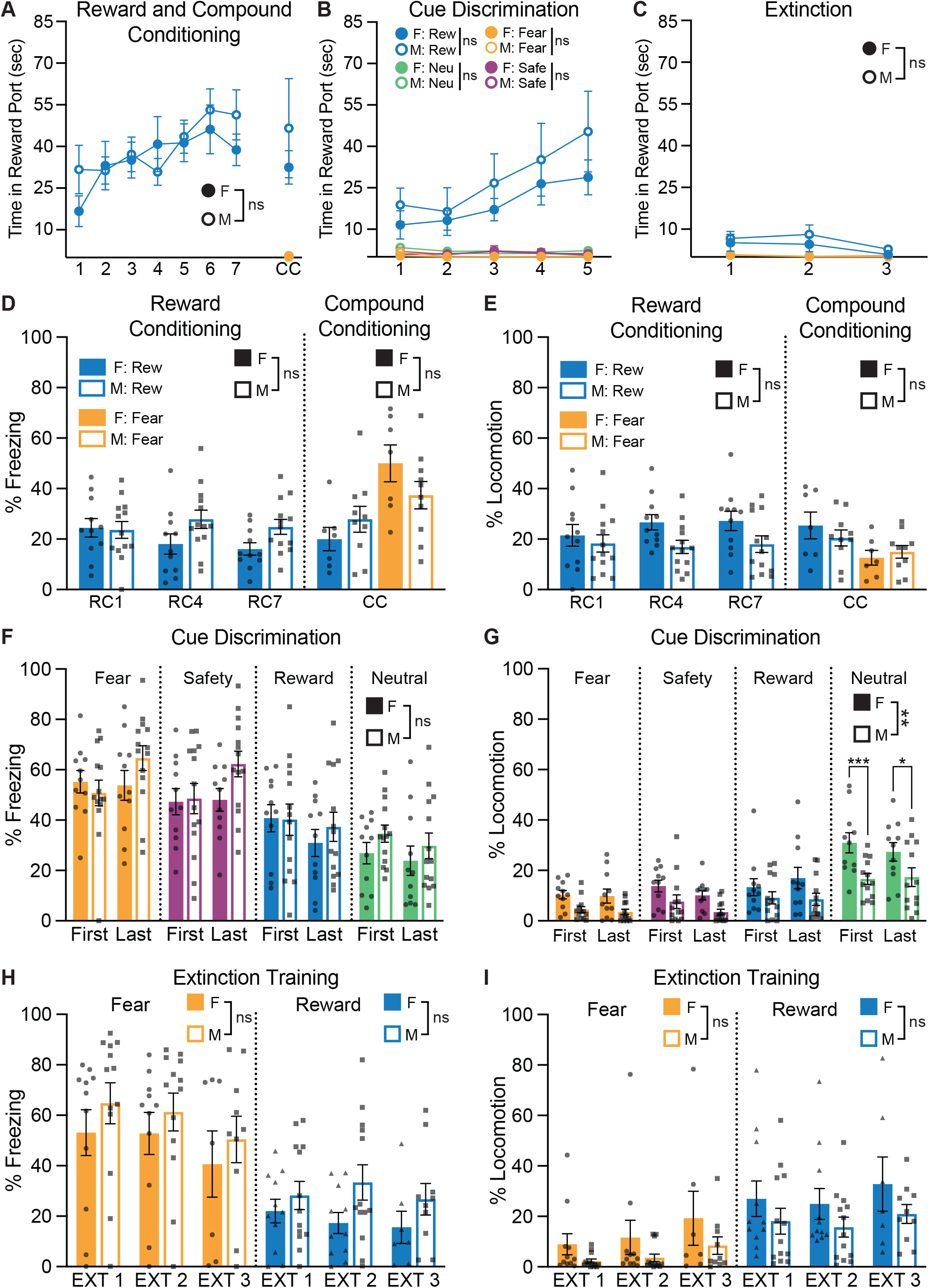
Reward and threat-related behaviors do not systematically vary by sex. A) Time spent in the sucrose port split between females (n=11; filled) and males (n=13; open) across reward conditioning and compound conditioning. There was no sex difference in port entry behaviors during the reward cue presentations (F(1,22)=0.1426, p=0.7093) nor the fear cue presentations (t(15)=1.770, p=0.0970) in early training phases. B) Time spent in the sucrose port for all cue types during cue discrimination training for females (filled circles) and males (open circles). There were no differences in time spent in port during reward cue (F(1,22)=0.6680, p=0.4225), fear cue (F(1,22)=1.808, p=0.1816), safety cue (F(1,22)=0.03567, p=0.8519), or neutral cue (F(1,22)=3.778, p=0.0648) presentations. C) During extinction training, females and males spent equivalent time in the sucrose port during reward cue (F(1,22)=0.2858, p=0.5983) and fear cue (F(1,22)=0.4472, p=0.5106) presentations. D) There were no significant main effects of sex (F(1,22)=1.1952, p=0.1763) or session (F(2,44)=1.818, p=0.1744) for percent of time freezing during cue presentations during reward conditioning, however the pattern of freezing slightly differed between sexes across the sessions: freezing decreased for the females, but stayed slightly elevated for the males (session x sex interaction, F(2,44)=4.566, p=0.0158). During compound conditioning, freezing was modulated by cue type (F(1,15)=35.79, p<0.0001), such that, on average, the fear cue elicited more freezing than the reward cue. The reward and fear cues prompted mildly different levels of freezing between females and males (cue type x sex interaction, F(1,15)=9.591, p=0.0074), but there was not a significant main effect of sex (F(1,15)=0.1031, p=0.7526). E) Percent of time locomoting during cue presentations was not significantly different between the sexes for either cue type during reward conditioning (F(1,22)=2.850, p=0.1055) and compound conditioning (F(1,15)=0.0984, p=0.7580). During compound conditioning, locomotion was modulated by cue type (F(1,15)=11.95, p=0.0035), and on average, the reward cue prompted more locomotion than the fear cue. F) There were no significant differences in freezing behavior between males and females for any cue type during the first (F(1,22)=0.07566, p=0.7858) or the last (F(1,22)=2.649, p=0.1179) day of the cue discrimination phase. G) When comparing locomotion to cue types during the first cue discrimination session, locomotion was modulated by cue type (F(3,66)=22.22, p<0.0001) and sex (F(1,22)=9.609, p=0.0052), especially during the neutral cue, where females locomote more during these presentations (p=0.0002). Locomotion during the final cue discrimination session was also modulated by cue type (F(3,66)=21.21, p<0.0001) and sex (F(1,22)=7.919, p=0.0099), specifically during the neutral cues (p=0.0478), where females locomote more during these presentations. H) Cue-evoked freezing behaviors did not differ between females and males during extinction training during fear cue presentations (F(1,22)=0.6988, p=0.4122) or reward cue presentations (F(1,22)=2.567, p=0.1233). However, the different cue types prompted different levels of freezing between females and males throughout extinction, where reward cue-induced freezing decreased in females, but was maintained in males (session x sex interaction, F(2,37)=4.852, p=0.0135). I) Cue-evoked locomotion behaviors did not differ between females and males during extinction training during fear cue (F(1,22)=1.700, p=0.2058) or reward cue (F(1,22)=1.581, p=0.2218) presentations. However, locomotion to the fear cue did increase across extinction training sessions (F(2,37)=5.376, p=0.0089). Data represent mean ± SEM. *p<0.05, *p<0.01, ***p<0.001, ****p<0.0001.

**Fig S3.**
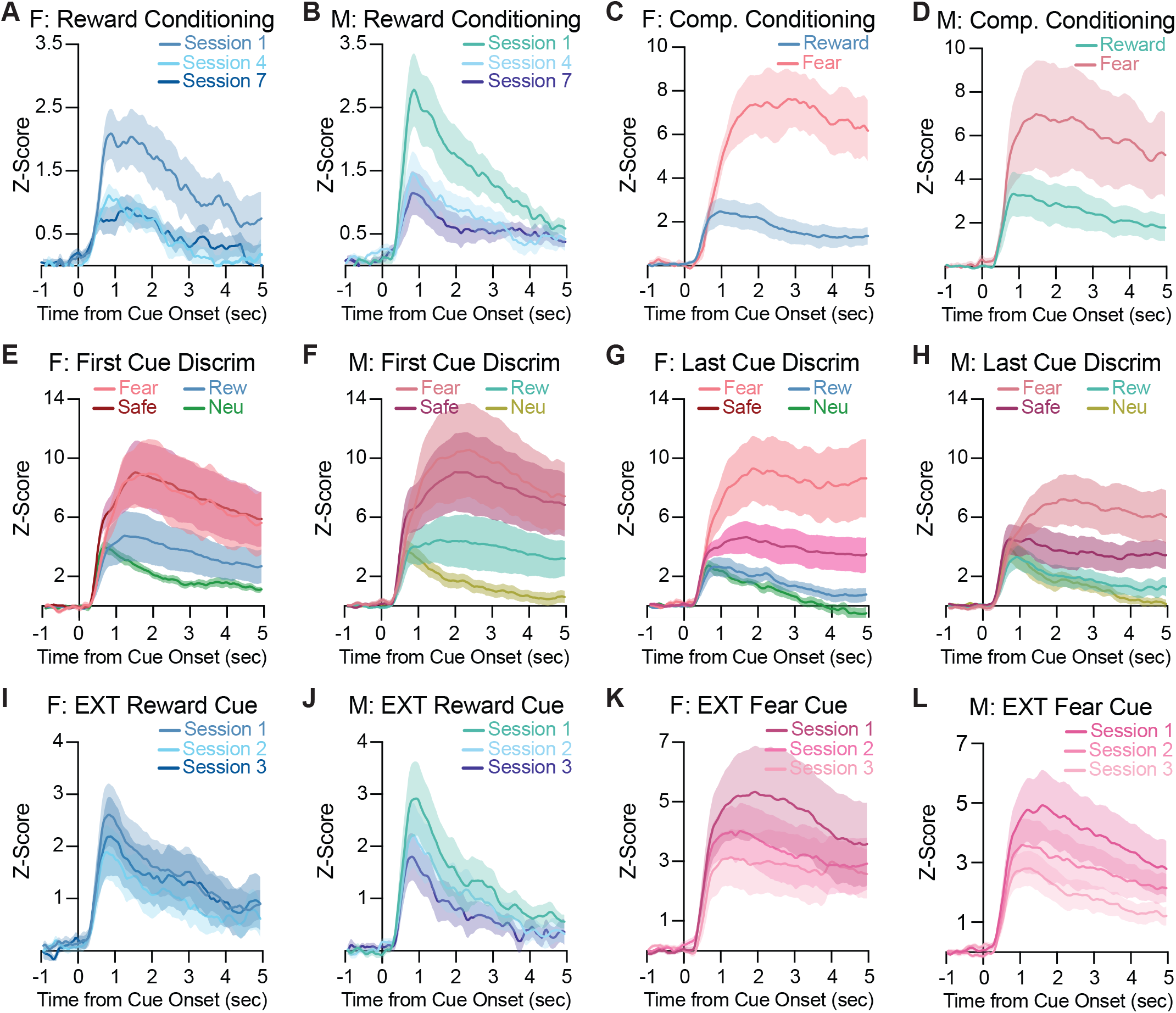
Cue evoked BLA dopamine transmission patterns or magnitudes do not systematically vary by sex. A) Female (n=11) BLA dopamine signals and B) male (n=13) BLA dopamine signals in response to reward cues during reward conditioning both featured moderate peaks that decreased across training. C) Female and D) male BLA dopamine dynamics during compound conditioning were characterized by large, sustained responses to the fear cue and modest responses to the reward cue. E) The BLA dopamine system responded smilarly in females and F) males during the first cue discrimination session. With higher dopamine transmission to the fear and safety cues, intermediate dopamine transmission to the reward cue, and lowest dopamine transmission to the neutral cue. G) By the last cue discrimination session, both females and H) males had the largest dopamine signals after the fear cue, intermediate dopamine transmission to the safety cue, and the lowest dopamine transmission to the reward and neutral cues. I) Reward cues during extinction training elicited moderate BLA dopamine dynamics in both females and J) males. K) Fear cues during extinction prompted relatively large dopamine signals that diminished with training in both females and K) males. Data represent mean ± SEM. F=female, M=male.

**Fig S4.**
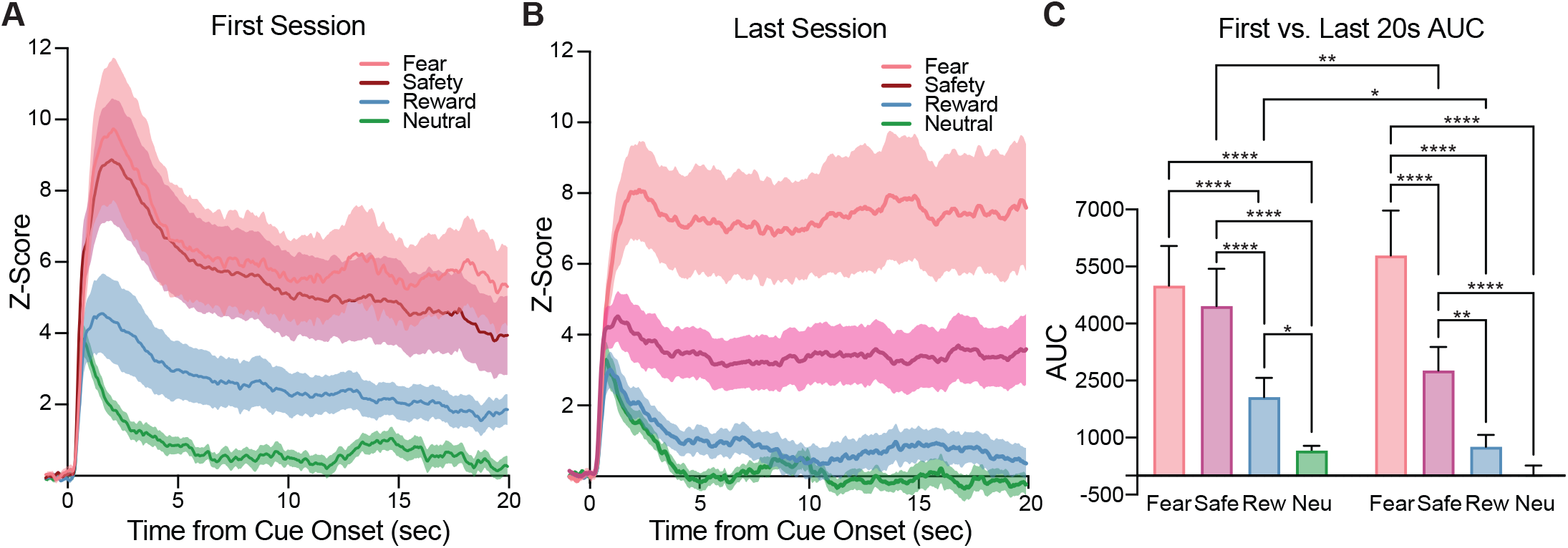
Fear and safety cues prompt larger and more sustained dopamine compared to the reward and neutral cues in both males and females during cue discrimination. A) In rats (n=24), on the first discrimination session all four cue types elicited a positive peak and remained above baseline for the full 20s cue presentation. Specifically, the fear and safety cues prompted larger, more sustained BLA dopamine traces compared to the reward and neutral cues. B) During the last discrimination session, BLA dopamine release remained highly sustained for the full 20s fear cue periods, while the dopamine release to the reward and neutral cues returned to baseline around 10s and 5s, respectively. The safety cue BLA dopamine trace had an intermediate magnitude and remained moderately sustained throughout the entire 20s presentation. C) AUC measures of the BLA dopamine signals show cue-induced dopamine was modulated by cue type (F(3,69)=21.96, p<0.0001) and this modulation changed across sessions (session x cue type interaction F(3,69)=4.562, p=0.0057). In the first discrimination session the fear cue trace is significantly larger than the reward (p<0.0001) cue trace, and the safety cue trace is also significantly larger than the reward (p<0.0001) cue trace. The neutral cue trace is significantly smaller than the fear (p<0.0001) cue trace, the safety (p<0.0001) cue trace, as well as the reward (p=0.0368) cue trace. The amounts of safety cue evoked dopamine (p=0.0015) and the reward cue evoked dopamine (p=0.0127) significantly decreased between the first and last cue discrimination sessions. On the last discrimination session, the fear cue elicited significantly larger dopamine transmission compared to all other cue types (safety: p<0.0001; reward: p<0.0001; neutral: p<0.0001), and the safety cue prompted significantly larger dopamine transmission compared to the reward (p=0.0012) and neutral (p<0.0001) cue types. Reward and neutral cue presentations evoked similar levels of BLA dopamine (p=0.4317). Data represent mean ± SEM. *p<0.05, **p<0.01, ***p<0.001, ****p<0.0001.

## Notes

### Competing Interest Statement

The authors have declared no competing interest.

### Summary of Updates

1. All datasets have been expanded with additional subjects. The whole paper has been revised to reflect some updated conclusions. 2. New analyses correlating dopamine cue responses with behavior across multiple experiments. Collectively, these analyses show that cue-evoked dopamine signals in the BLA, for either rewards or threats, do not reliably correlate with cue-evoked behavioral responses (reward seeking or freezing). 3. New analyses showing within-session time courses of cue discrimination in dopamine signals. These show that dopamine signals differentiate cue types across the initial discrimination session, suggesting that these signals are a dynamic read out of relative stimulus salience. 4. New analyses comparing the reward cue-evoked signals across experimental phases (New Fig 5). This shows that BLA dopamine signals can dynamically scale based on the emotional uncertainty of the learning environment. 5. A new appetitive valence scaling experiment has been added comparing BLA dopamine signals for sucrose versus a higher value reward, Ensure (New Fig 2). These new data show that despite rats preferring Ensure reward behaviorally, the evoked dopamine signals are the same as for sucrose. 6. A new aversive valence scaling experiment comparing BLA dopamine signals to low vs moderate footshocks. These data show, in contrast to the stable reward scale, BLA dopamine increases as aversive stimulus intensity increases. 7. A new threat scaling experiment (New Figure 7) where rats discriminate between cues predicting low vs moderate footshocks. This shows that the cue-evoked BLA dopamine signal scales with threat intensity. 8. A new optogenetic activation experiment where VTA dopamine terminals in the BLA were excited on a subset of fear cue presentations. These data show that augmenting dopamine in the BLA does not acutely alter expression of freezing behavior.

